# Jade1 and the HBO1 histone acetyltransferase complex are spatial-selective cofactors of the pluripotency transcription factor Oct4

**DOI:** 10.1101/2024.11.07.622531

**Authors:** Yifan Wu, Asit K. Manna, Li Li, Zuolian Shen, Hiroshi Handa, Mahesh B. Chandrasekharan, Dean Tantin

## Abstract

Oct4 is a master regulator of pluripotency. Potential Oct4 interactors have been cataloged but the manner and significance of these interactions are incompletely defined. Oct4 is capable of binding to DNA in multiple configurations, however the relationship between these configurations and cofactor recruitment (and hence transcription output) are unknown. Here, we show that Oct4 interacts with common and unique proteins when bound to DNA in different configurations. A unique protein is Jade1, a component of HBO histone acetyltransferase complexes. Jade1 preferentially associates with Oct4 when bound to <u>M</u>ore palindromic <u>O</u>ctamer-<u>R</u>elated <u>E</u>lement (MORE) DNA sequences that bind Oct4 dimers. Surprisingly, we find that the Oct4 N-terminal activation domain, rather than facilitating Jade1 binding, serves as an autoinhibitory domain that dampens the interaction. ChIP-seq using HBO1, the enzymatic component of the complex, identifies a preference for binding adjacent to Oct4 at MORE sites. Using purified recombinant proteins and nucleosome complexes, we show that the HBO1 complex acetylates histone H3K9 within nucleosomes more efficiently when Oct4 is co-bound to a MORE. An Oct4 mutant with superior MORE binding characteristics also shows superior ability to catalyze H3K9 acetylation. Jade1 knockdown reduces H3K9Ac at regions where Oct4 binds a MORE but not a simple octamer. Cryo-electron microscopy reveals that Oct4 bound to a MORE near the nucleosome entry/exit site partially unwinds DNA from nucleosome core particles, and identifies additional mass associated with the HBO1 complex. These results identify a novel mechanism of transcriptional regulation by Oct4.

## Introduction

Gene transcription is regulated by the expression and activity of sequence specific DNA binding transcription factors (1). Transcription factors regulate different target genes through recruitment of transcription cofactors and action on local chromatin. For example, chromatin modifying enzymes such as histone acetyltransferases (2–5) can interact with transcription factors, enhance local histone acetylation and favor transcription. Although the binding of a given transcription factor to different target sequences within the genome is known to be modulated by interactions with other co-expressed transcription factors (6), whether the configuration of transcription factor interactions with DNA can influence their interactions with cofactors and hence transcription output is under-explored.

Octamer binding (Oct)/Pit-Oct-UNC (POU) proteins are a group of extended-family homeodomain proteins that play varied roles in development and reprogramming, stress response, somatic stem cell function and immune function (7, 8). The canonical monomeric binding site used by class II and class V POU proteins (including the pluripotency regulator Oct4/Pou5f1) is the octamer motif, 5’ATGCAAAT (9). Variant DNA binding sites have also been described, including two dimeric binding sites known as a PORE (palindromic octamer related element) and a MORE (10, 11).

Structures of the POU proteins Oct1 and Oct4 bound to DNA in different configurations, including bound to canonical octamer motifs, MOREs and POREs, have been determined by X-ray crystallography and cryo-electron microscopy (cryoEM) (12–15). These studies reveal that while Oct proteins bound to a PORE resemble a dimeric arrangement of two adjacent monomers, binding to a MORE induces a rearrangement of the two DNA binding subdomains (termed POUS and POUH) such that both subdomains of the same protein bind to adjacent major grooves on one side the DNA helix, rather than spanning the DNA as in a canonical octamer and a PORE. This unique binding configuration generates unique chemical surfaces of the DNA binding domains and in the orientations of the N- and C-termini, with the potential to recruit distinct transcriptional coregulators. Oct1 and Oct4 binding to MOREs can be induced by oxidative stress through phosphorylation of the POUH DNA binding subdomain, which favors intramolecular hydrogen bonding between the DNA binding subdomains to stabilize MORE binding (16). The stabilization can be made constitutive with a phospho-mimetic mutation (16). This study also identified MOREs in the promoter regions of widely expressed genes such as *Ell*, *Polr2a*, *Rras* and *Rras2*.

Here, we identify Jade1 as a selective interactor of Oct4 bound to MORE-containing but not octamer-containing DNA. Jade1 is a component of histone acetyltransferase complexes that contain the MYST family histone acetyltransferase (HAT) HBO1/KAT7/MYST2 as the catalytic subunit.

Deleting Oct4’s N-terminal activation domain augments the interaction with Jade1. We show that HBO1 and Oct4 are preferentially co-enriched at MOREs in mouse pluripotent embryonic stem cells (ESCs), and that Jade1/HBO1 complexes more robustly acetylate H3K9 when Oct4 is supplied to nucleosomes containing a MORE near the entry/exit site in a dose-dependent fashion. A mutant form of Oct4 that binds to MOREs with superior affinity is also superior at catalyzing H3K9 acetylation.

Jade1 knockdown in mouse ESCs decreases H3K9Ac levels at the MORE region near *Polr2a* but not the compound Oct:Sox site in a *Pou5f1* enhancer. Finally, we use cryoEM to show that Oct4 bound to MOREs near the nucleosome entry/exit site are associated with DNA unwinding, and identify additional densities associated with the HBO1/Jade1L complex surrounding the nucleosome core particle. These studies identify Jade1 as a spatioselective cofactor of Oct4.

## Results

### Oct4 and its cofactors can be enriched by magnetic beads coupled to simple octamer or MORE DNA

To identify interacting proteins with potential selectivity for Oct4 bound to DNA in different configurations, we used magnetic Ferrite-Glycidyl methacrylate (FG) beads covalently attached to double-stranded DNA containing either multimerized octamer elements or MORE sequences. In both cases, point mutant DNA was used as a control (Fig. 1*A*). To reduce the effect of Oct1, an Oct4 paralog co-expressed in ESCs, we enriched Oct4 from Oct1 deficient ESC extracts (17).

**Figure 1.**
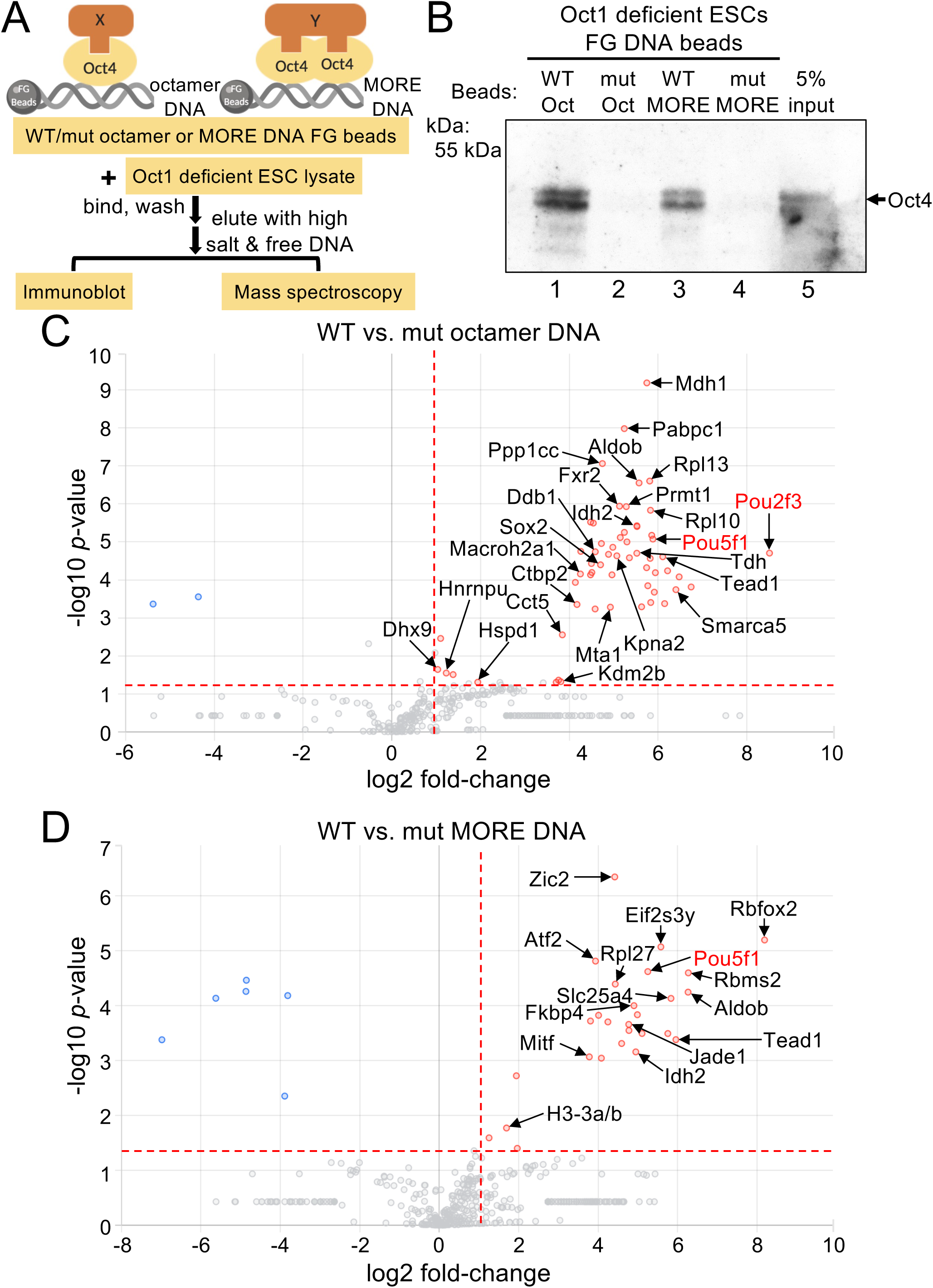
Oct4 DNA affinity purification identifies Jade1 as a spatioselective interaction partner. *A*, Bead affinity capture workflow. Oct1 deficient ES cell lysates were incubated wild-type or mutant octamer or MORE DNA sequences immobilized on FG beads. Proteins were eluted using high salt and free excess DNA, the presence of Oct4 in the eluates verified by Western Blot, and the presence of associated proteins analyzed by mass spectrometry. *B*, Material eluted from (A) was immunoblotted using Oct4 antibodies as a quality control. *C*, MS-Mascot output showing the 58 proteins enriched using wild-type octamer relative to mutant DNA. Average fold enrichment vs. negative inverse log10 *p*-value is shown based on N=3 independent experiments. Threshold lines show 2-fold enrichment and *p*=.05. Black text indicates proteins identified in prior studies. Red text shows POU proteins. Octamer-binding POU transcription factor proteins are shown in red. *D*, Similar analysis using MORE DNA. 27 proteins were significantly enriched.

Undifferentiated Oct1-deficient ESCs are hypersensitive to oxidative stress but otherwise normal in terms of gene expression, morphology, and immortality (17, 18). Oct4 bound to wild-type but not mutant DNA sequences (Fig. 1*B*). The co-enriched materials were then analyzed by ultra-high mass range hybrid quadrupole-orbitrap mass spectrometry (Table S1). Comparing material enriched using wild-type vs. mutant octamer DNA (≥2-fold change, *p*<0.05), there were 58 proteins significantly enriched using wild-type DNA, while only 2 proteins were enriched with mutant (Fig. 1*C*). Proteins enriched using wild-type octamer sequence DNA included Pou5f1 (Oct4), Pou2f3 (Oct11) and Sox2, the latter of which is known to pair with Oct4 at Oct-Sox motifs (14). 15 other proteins were previously identified in different reports of Oct4 cofactors (Fig. 1*C*, black text). Similarly, comparing material enriched using beads coupled to wild-type vs. mutant MORE DNA, there were 27 proteins enriched using wild-type DNA pull-down vs. 6 enriched with point mutant MORE DNA. Proteins enriched using wild-type MORE DNA included once again Pou5f1 (Oct4) and 7 other proteins known to interact with Oct4 (Fig. 1*D*, black text). These results support the reliability of the mass spectrometry findings.

Comparing the wild-type octamer and MORE enriched proteins, 89.5% (51 out of 57) and 76.9% (20 out of 26) of Oct4 cofactors were specific to either the octamer or MORE sequence DNA. Other than Oct4 itself, only seven proteins overlap (Fig. S1*A*). This finding is consistent with a hypothesis that different Oct4 configurations preferentially recruit different cofactors. Comparing the enriched proteins with three prior reports on Oct4 interaction partners (19–21), we identified 16 overlapping proteins (Fig. S1*B*).

### Jade1 associates with Oct4 bound specifically to MORE-containing DNA

One protein strongly co-enriched with Oct4 using beads coupled to wild-type MORE DNA was Jade1, a PHD finger-containing protein thought to act as an adapter subunit for the HBO1 HAT complex.

Jade1 contacts histone H3 and H4 to facilitate HBO1 complex binding to chromatin (22–24). This complex is reported to acetylate histones H3 and H4 to activate transcription (25). To test if Jade1 acts as an Oct4 cofactor specifically at MORE sequences, we immunoblotted for Jade1 enriched from Oct1 deficient mouse ESC extracts and FG beads coupled to the four types of DNA sequences (wild-type or mutant octamer or MORE). Only material enriched using wild-type MORE-coupled beads showed enrichment for Jade1 (Fig. 2*A*). We then performed co-immunoprecipitation (co-IP) experiments using Oct1 deficient ESC extracts and antibodies against Oct4 and Jade1. Both co-IP using Oct4 antibodies and Jade1 Western blot, and reciprocal Jade1 co-IP with Oct4 Western blot identified specific bands (Fig. 2*B* and *C*). This interaction was confirmed using extracts from wild-type ESCs (Fig. S1*C*), indicating that the interaction was not an artifact induced by the absence of bound Oct1.

**Figure 2.**
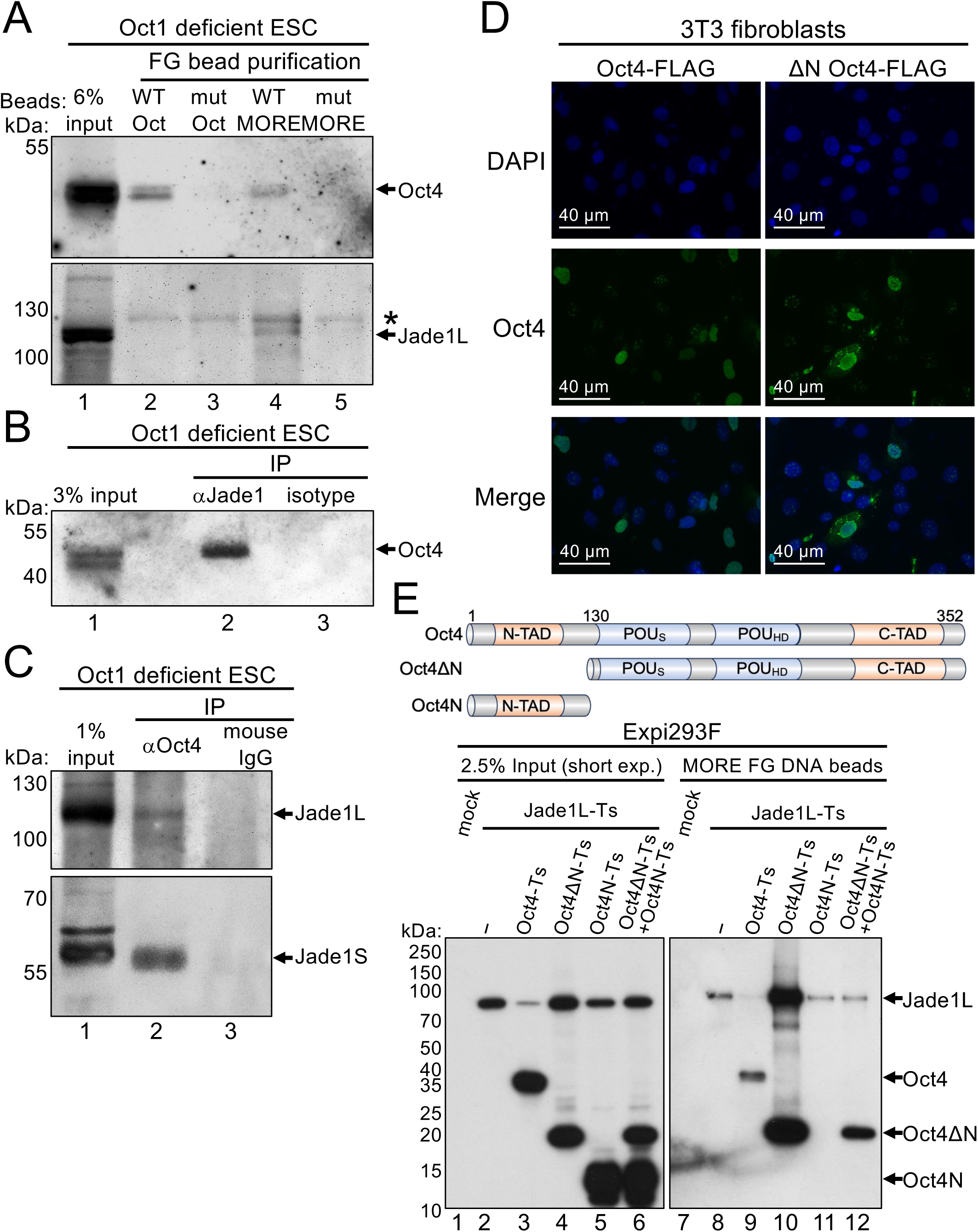
Oct4 interacts with Jade1. *A*, Oct4 and associated proteins were precipitated from Oct1 deficient ESCs using FG beads immobilized with wild-type or mutant octamer and MORE DNA sequences. The isolated material was immunoblotted using anti-Oct4 and anti-Jade1 antibodies. *B*, Extracts from Oct1 deficient ESCs were used for immunoprecipitation with anti-Jade1 antibodies, or corresponding rabbit IgG as a negative control, and immunoblotted with anti-Oct4. *C*, Oct1 deficient ESCs were immunoprecipitated using anti-Oct4 antibodies or mouse IgG as a negative control, and immunoblotted with anti-Jade1. *D*, 3T3 cells were transiently transfected with plasmids encoding FLAG- and twin-strep-tagged Oct4 or ΔN Oct4. After 48 hr, cells fixed and stained with antibodies against Oct4 (Donkey anti-mouse Oct4-Alexa 488) and imaged using an EVOS M7000 microscope. *E*, Expi293F cells were transiently co-transfected with plasmids encoding FLAG- and twin-strep-tagged Oct4, ΔN Oct4 or Oct4’s N-terminus, and twin-strep-tagged Jade1L. After 48 hr, lysates were prepared in the presence of MG-132 and incubated with MORE DNA-coupled FG beads, washed, resolved using SDS-PAGE, and probed with HRP-conjugated anti-StrepII antibodies.

### The Oct4 N-terminus dampens Jade1 recruitment to MORE-containing DNA

Oct4’s N-terminus has been shown to harbor an activation domain (26). To test if this domain is required for the interaction with Jade1, we deleted Oct4’s N-terminal 130 amino acids, preserving the DNA binding domain. Prior work has also shown that the Oct4 DNA binding domain contains a nuclear localization sequence (27), suggesting that an N-terminally truncated protein retaining the DNA binding domain should still localize to nuclei. We confirmed this prediction using immunofluorescence in 3T3 fibroblasts and antibodies against Oct4 (Fig. 2*D*). We then performed FG bead MORE DNA affinity purification using Expi293F cells transfected with plasmid constructs expressing either full-length or N-terminally deleted Oct4, as well as the large Jade1 isoform Jade1L. All constructs contained twin-strep tags for efficient detection. N-terminally truncated Oct4 purified from Expi293F cells was selectively degraded during incubation with FG beads, but this could be blocked by including the proteasome inhibitor MG-132 in the lysates (not shown). This experiment was therefore conducted using 20 μg/mL MG-132 in all lysates. Full-length and N-terminally truncated Oct4 differentially affected co-expressed input Jade1L protein levels, with full-length Oct4 significantly reducing Jade1L levels and N-terminal deletion restoring levels to that of parent transfections lacking Oct4 (Fig. 2*E*, lanes 2-4). Expression of the Oct4 N-terminus alone or in conjunction with N-terminally deleted Oct4 was insufficient to reduce co-transfected Jade1, indicating that the N-terminus must be tethered to the Oct4 DNA binding domain (lanes 5-6). Jade1L showed a low level of baseline MORE bead binding in the absence of co-transfected Oct4 (lanes 7-8), either due to nonspecific affinity for DNA or the presence of one or more interacting proteins capable of binding these DNA sequences in the Expi293F lysates. Interestingly, Oct4 lacking its N-terminus more efficiency copurified Jade1 compared to wild-type (compare lane 10 to lane 9). Separately co-expressed Oct4 N-terminus was sufficient to reduce this augmented interaction (lanes 11-12). These results indicate that Oct4’s N-terminus inhibits the interaction with Jade1.

### HBO1 preferentially associates near MOREs in ESCs

We attempted to determine Jade1 binding preferences genome-wide using Jade1 ChIP-seq (chromatin immunoprecipitation and sequencing) with an anti-Jade1 antibody without success. We then turned to ChIP using antibodies against the catalytic subunit of the Jade1/HBO1 complex, HBO1, which has been used successfully in ChIP-seq with HeLa cells (25). Approximately 13,800 peaks were identified with high confidence (Table S2). HBO1 showed a preference for promoter binding, with 37% of the HBO1 ChIP peaks within ±1 kb of transcription start sites (TSS, Fig. S2*A*), broadly consistent with findings from HeLa cells (25). By aligning peaks with TSSs, we found that HBO1 is most highly enriched on either side of the TSS (Fig. S2*B*). We then compared the HBO1 enrichments to Oct4 ChIP-seq data also generated using mouse ESCs (28). A significant fraction of HBO1 ChIP-seq peaks (1744) overlapped (<200 bp) with Oct4 peaks (Fig. 3*A*). Studying individual genome tracks, we found that HBO1 ChIP peaks either overlapped with or were immediately adjacent to MORE containing genes such as *Polr2a*, *Zmiz2* and *Rras2*, but were not identified within 2 kb of octamer containing genes such as *Pou5f1* (Fig. 3*B*). To quantify this more precisely, we calculated the average distance between Oct4 and the nearest HBO1 peak in the total dataset vs. the distance between Oct4 bound at a MORE and the nearest HBO1 peak. The distance shifted from >10 kb to <200 bp (Fig. S2*C*, Table S3).

**Figure 3.**
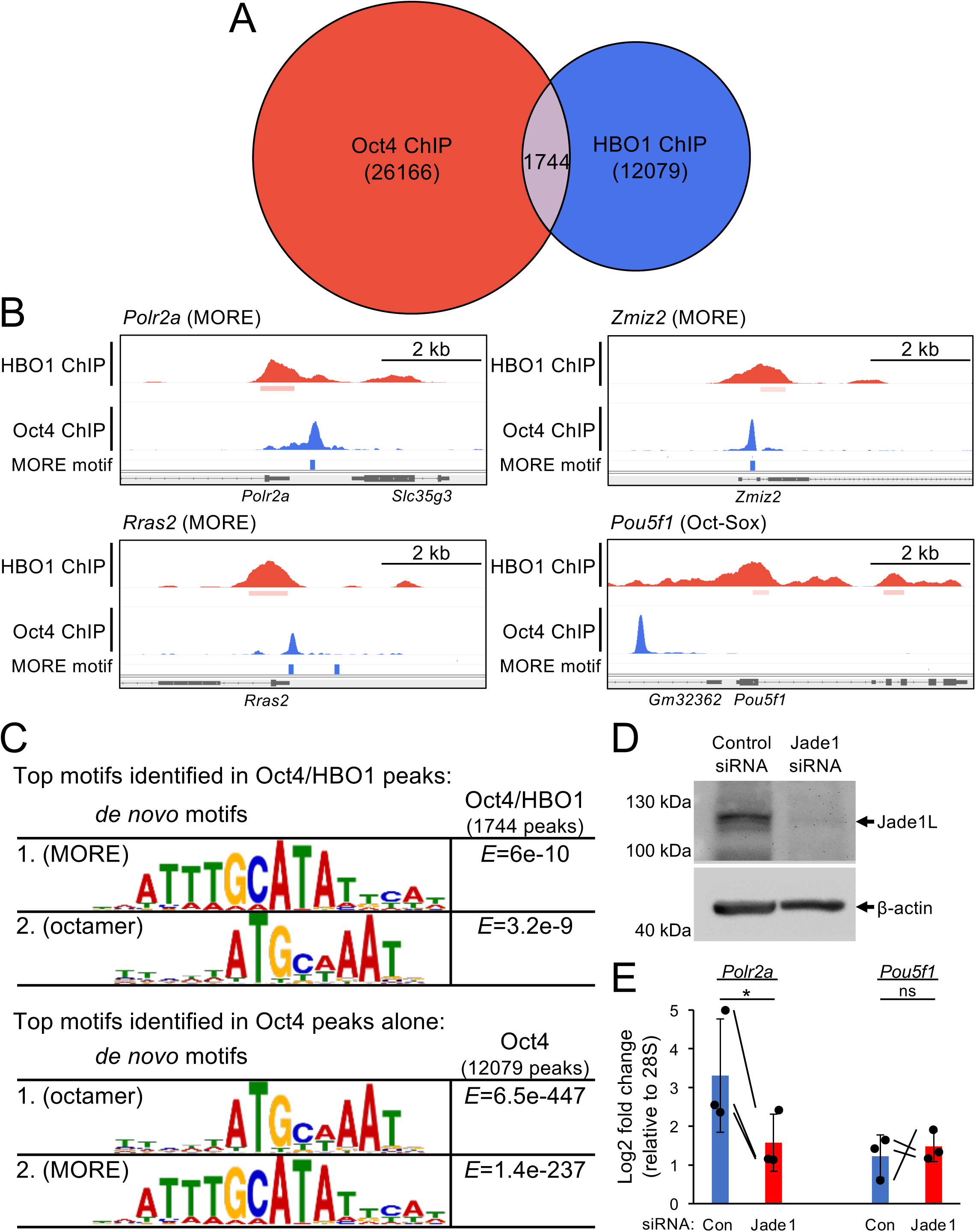
Jade1/HBO1 complex prefer binding at MORE sites compared to Octamer sites. *A*, Venn diagram showing overlap between Oct4 and HBO1 ChIP-seq peaks. *B*, HBO1 and Oct4 ChIP-seq peaks at example MOREs (*Polr2a*, *Zmiz2* and *Rras2*) and an octamer site (*Pou5f1*). *C*, Motif identification using MEME-ChIP at Oct4 ChIP-seq overlapping with HBO1 (≤100 bp between peak centers) vs. all Oct4 ChIP-seq peaks. *D*, Immunoblot showing Jade1L levels 48 hr post-transfection of mouse ESCs with control or Jade1-specific siRNAs. β-actin is shown as a loading control. *E*, Jade1 knockdown and control ESCs treated similarly to (*D*) were subjected to ChIP-qPCR using anti H3K9Ac antibodies and primer-pairs spanning two adjacent MOREs upstream of *Polr2a*, spanning the Oct:Sox site in the *Pou5f1* upstream enhancer, or a 28S ribosomal RNA control region. Enrichment is shown relative to 28S, which was set to 1.0. N=3 biological replicates. Significance was ascribed using paired two-tailed T-tests. Error bars denote ±standard deviation.

We used MEME-ChIP to directly identify sequences enriched in Oct4 peaks alone compared to overlapping Oct4/HBO1 peaks. Oct4/HBO1 co-enriched targets identified by ChIP-seq preferentially show enrichment of the MORE over the octamer motif, while Oct4 peaks in isolation, though more abundant, enriched octamer motifs more robustly than MOREs (Fig. 3*C*). These data indicate that the HBO1/Jade1 complex is recruited by Oct4 more frequently to the MOREs compared to octamer sites.

To further test the relationship between Oct4 MORE binding and H3K9 acetylation mediated by Jade1, we combined transient knockdown of Jade1 in mouse ESCs with H3K9Ac ChIP-qPCR using MORE-containing and simple octamer-containing regions. Prior work has established that Jade1 loss of function by gene trap is compatible with development (29). We transfected control and Jade1-specific small interfering RNAs (siRNAs) into mouse ESCs, resulting in efficient Jade1 knockdown (Fig. 3*D*). Similar cells were then used in ChIP-qPCR experiments using H3K9Ac antibodies and qPCR primers specific to *Por2a* (MORE), *Pou5f1* (octamer) and 28S (nonspecific normalization control). Relative to 28S, Jade1 knockdown resulted in lower levels of H3K9Ac levels at the MORE-containing *Polr2a* gene (Fig. 3*E*). These findings indicate that H3K9 acetylation levels near an Oct4-bound MORE are dependent on Jade1.

### Purified HBO1/Jade1L complexes acetylate histone lysines within nucleosomes

We obtained mammalian expression constructs for recombinant twin-strep-tagged Jade1L (30) and purified the protein from Expi293F cells using similar methods as for Oct4. We also subcloned Jade1S from this construct allowing us to purify the smaller isoform. Analysis of the purified Jade1L and Jade1S proteins by SDS-PAGE and Coomassie staining revealed the presence of Jade1L and Jade1S at ∼100 and ∼50 kDa, respectively, as well as several additional bands (Fig. S3*A*), which were identified by immunoblotting as HBO-1 (∼75 kDa), ING4 (∼35 kDa) and hEaf6 (∼25 kDa, Fig. S3*B*). Because the purified Jade1L complex contains the catalytic HBO1 activity and other associated components, we hypothesized that the purified material would acetylate one or more histone lysines within nucleosomes in vitro. The Jade1L/HBO1 complex is known to acetylate histone H4, while the alternative complex of HBO1 with BRPF1 is known to acetylate histone H3K14 (22, 31). We reconstituted nucleosome complexes in the presence of the HBO1/Jade1/ING4/hEaf6 complex and Oct4 using purified recombinant proteins (Fig. S4) and nucleosomal DNA containing a MORE (Fig. 4*A*) incubated at equimolar ratios for 30 min over ice. We confirmed that purified Jade1L complex can efficiently acetylate histone H4 within nucleosomes (Fig. 4*B*, lane 2). Blotting with specific antibodies indicates that the purified material can acetylate H4K5, H4K8 and H4K12 but not H4K16 (Fig. 4*C*, lane 2). Immunoblotting using antibodies specific to H3 acetylation revealed no acetylation on H3K14 as expected, but surprisingly acetylation at H3K9, though this antibody did show some cross-reactivity to unacetylated H3. H3K18 and H3K27 were also not acetylated by purified Jade1L/HBO1 complex (Fig. 4*C*, lane 2). A second H3K9Ac antibody showed the same specific acetylation but with background cross-reactivity (not shown). H3 and H4 acetylation events are quantified from triplicate biological replicate experiments in Fig. 4*D*.

**Figure 4.**
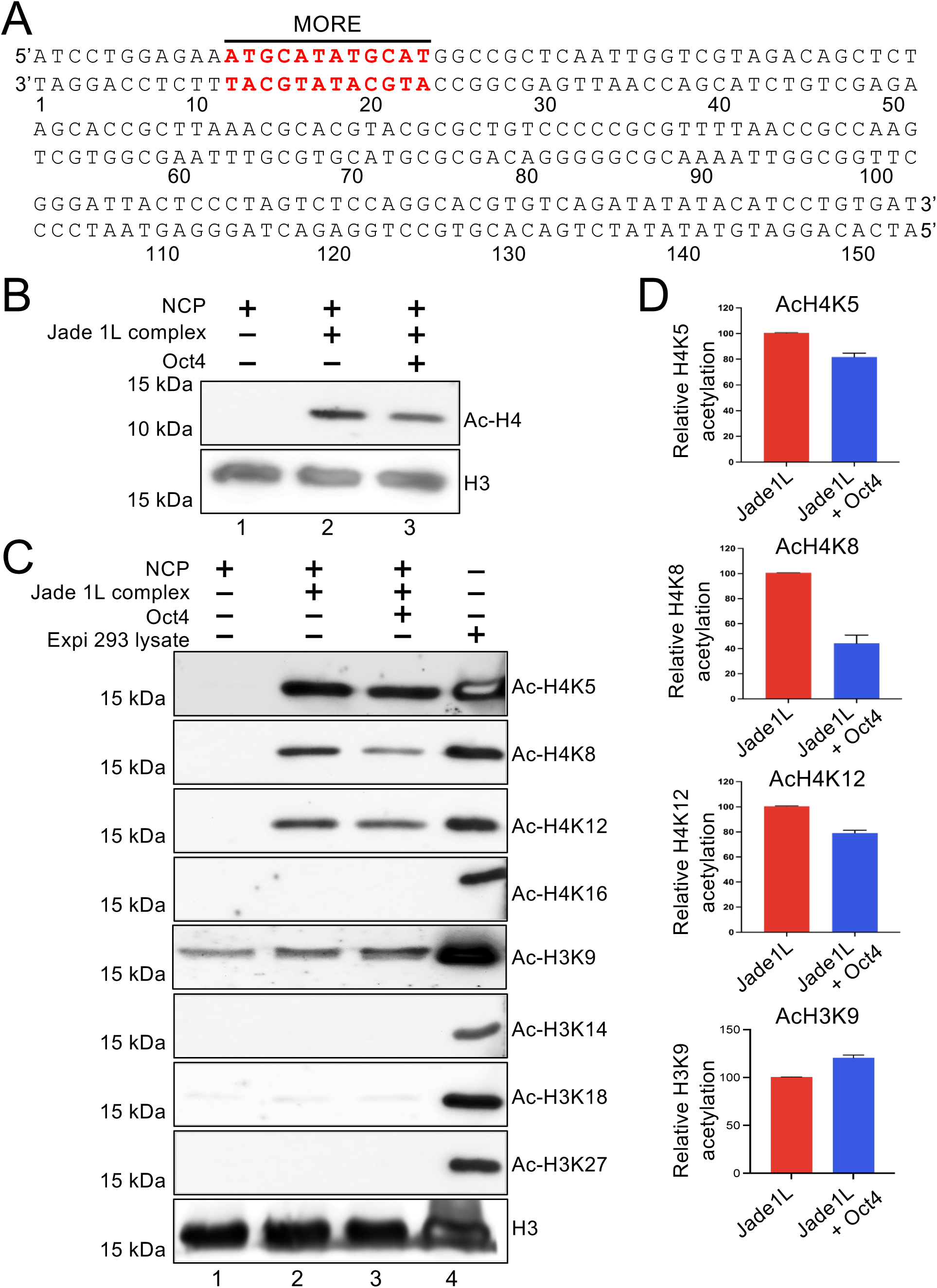
Oct4 augments acetylation of histone H3K9 by the HBO1/Jade1L complex. *A*, The DNA sequence used for reconstituting nucleosomes containing MORE sequence near the entry/exit site. The MORE sequence is indicated in red. *B*, HAT assay using nucleosome core particles (NCPs) containing a MORE. NCPs were incubated with purified Jade1L complex in the presence or absence of Oct4. NCP alone was included as a control (lane 1). Reactions were immunoblotted with antibodies against acetylated histone H4. Pan-H3 antibodies were included as a loading control. *C*, Acetylation of specific histone H4 and H3 tail lysines was detected by immunoblotting. *D*, Quantitation of acetylation of specific histone lysines by the HBO1/Jade1L complex in the absence or presence of Oct4. Quantitation was performed using Image J software from N=3 independent experiments. Error bars indicate standard deviation. Data were plotted using GraphPad Prism software.

### Oct4 bound to a MORE augments histone H3K9 acetylation by Jade1L/HBO1

We tested the effect of Oct4 bound to MORE DNA on Jade1L/HBO1-mediated in vitro HAT activity using purified nucleosomes as a substrate. Surprisingly, acetylation at all histone H4 lysines was reduced or unchanged upon inclusion of purified Oct4, whereas the acetylation on lysine 9 on histone H3 was increased (Fig. 4*C*, lane 3). To determine if there is any effect on Oct4 binding stoichiometry in presence of Jade1L, we tested Oct4 binding to MORE containing DNA in the presence and absence of the HBO1/Jade1/ING4/hEaf6 complex. In the presence of Jade1L, the proportion of dimeric DNA-bound Oct4 increased slightly (Fig. 5*A*, compare lane 4 to lane 2). Oct1 can also bind to MOREs (13), and point mutation of Oct1 at S385D on a loop region of POUH promotes intramolecular hydrogen bonding with POUS to stabilize the dimeric MORE-bound configuration (16). Because this region is highly conserved between Oct1 and Oct4, we generated the analogous mutation in the Oct4 DNA binding domain, Oct4^S229D^, in the context of the Oct4 expression construct (Fig. S5*A*). As expected, Oct4^S229D^ bound more efficiently to the MORE compared to Oct4 (Fig. 5*B*). Interestingly, Oct4^S229D^ augmented H3K9 acetylation by both Jade1L- and Jade1S-containing HBO1 HAT complexes more strongly compared to wild-type Oct4 (Fig. 5*C*-*D*). We also used increasing concentrations of mutant Oct4 to show a dose-dependency for catalysis by both Jade1L (Fig. 5*E*) and Jade1S (Fig. S5*B*). The H3K9 acetylation signal observed with no Jade 1/HBO1 complex was also observed when acetyl-CoA was omitted (not shown), indicating that the background signal is due to antibody cross-reactivity with unacetylated H3 (Fig. 5*E*, lane 1). Jade1L and Jade1S quantification from N=3 and N=4 independent experiments, normalized to H4, is shown in Fig. 5*F* and Fig. S5*C*.

**Figure 5.**
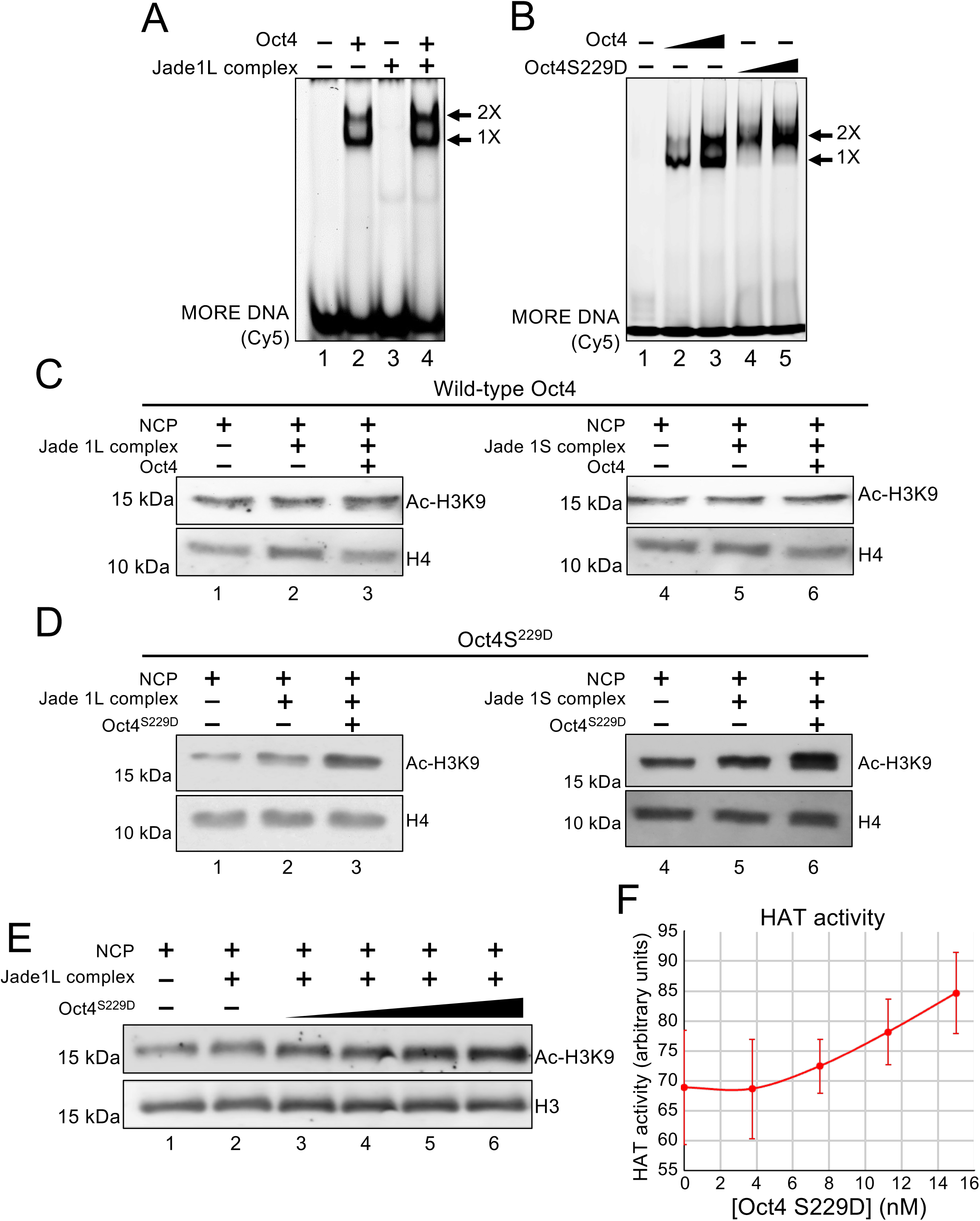
An Oct4 mutant that more efficiently binds to MOREs more efficiently acetylates nucleosomes in the presence of the HBO1/Jade1L complex. *A*, EMSA showing Oct4 protein binding to Cy5-labeled MORE DNA probe with or without the Jade1L/HBO1 HAT complex. Monomeric and dimeric interactions with the DNA are indicated by the arrows. *B*, Comparison of increasing concentrations of wild-type and S229D Oct4 binding to a MORE DNA probe. Monomeric and dimeric interactions bands are also shown by the arrow. *C*, Purified Jade1L and Jade1S containing components of the HBO1 HAT complex were used with wild-type Oct4 in HAT assays using nucleosome monomers containing MORE DNA. *D*, Similar data shown using mutant Oct4^S229D^. *E*, HAT assay using MORE-containing nucleosome core particles (NCPs) treated with 200 ng purified Jade1L complex in the presence of increasing amounts of purified recombinant Oct4. Histone H4 is shown as a loading control. The extent of histone H3K9 acetylation was detected by Western blotting. *F*, Quantification from N=4 experiments using the ratio of acetylated H3K9 to H4 as an internal standard. Error bars denote ±standard deviation.

### CryoEM structure of the HBO1/Jade1/ING4/hEaf6 complex and Oct4^S229D^ in the presence of a nucleosome

We reconstituted the HBO1/Jade1/ING4/hEaf6 nucleosome complex in the presence of Oct4^S229D^ using purified recombinant proteins (Fig. S3, S4) incubated at equimolar ratios for 30 min over ice. We processed 6200 dose-fractionated cryoEM images (Fig. 6*A* and Fig. S6*A*) to elucidate the structure of the complex. One group of 2D classifications (Fig.6*B*) was further refined to generate the best reconstructed and refined 3D volume, with 7.2 Å resolution and a B Factor of 322 (Fig. 6*C* and Fig. S6*A*, center). The refined volume showed an extension of DNA and extra volume surrounding the nucleosome core resulting in a more spherical structure (Fig. 6*C*), consistent with a nucleosome/HAT complex. The structural resolution was insufficient to model the position of HBO1, Jade1, ING4 or hEaf6, however the dimeric Oct4 POUS and POUH domains (PDB ID: 7XRC) could be manually fitted into the extended volume (Fig. 6*D*, left and right). A nucleosome core structure (PDB ID: 6TEM) could also be fitted into the 3D volume, with the 5’ part of the DNA sequence removed and replaced with linear B-form DNA to complete the volume (Fig. 6*E*). This result is consistent with an interpretation in which dimeric Oct4 bound near the nucleosome entry/exit site in the presence of the HBO1 HAT complex partially unwinds DNA from core nucleosomes.

**Figure 6.**
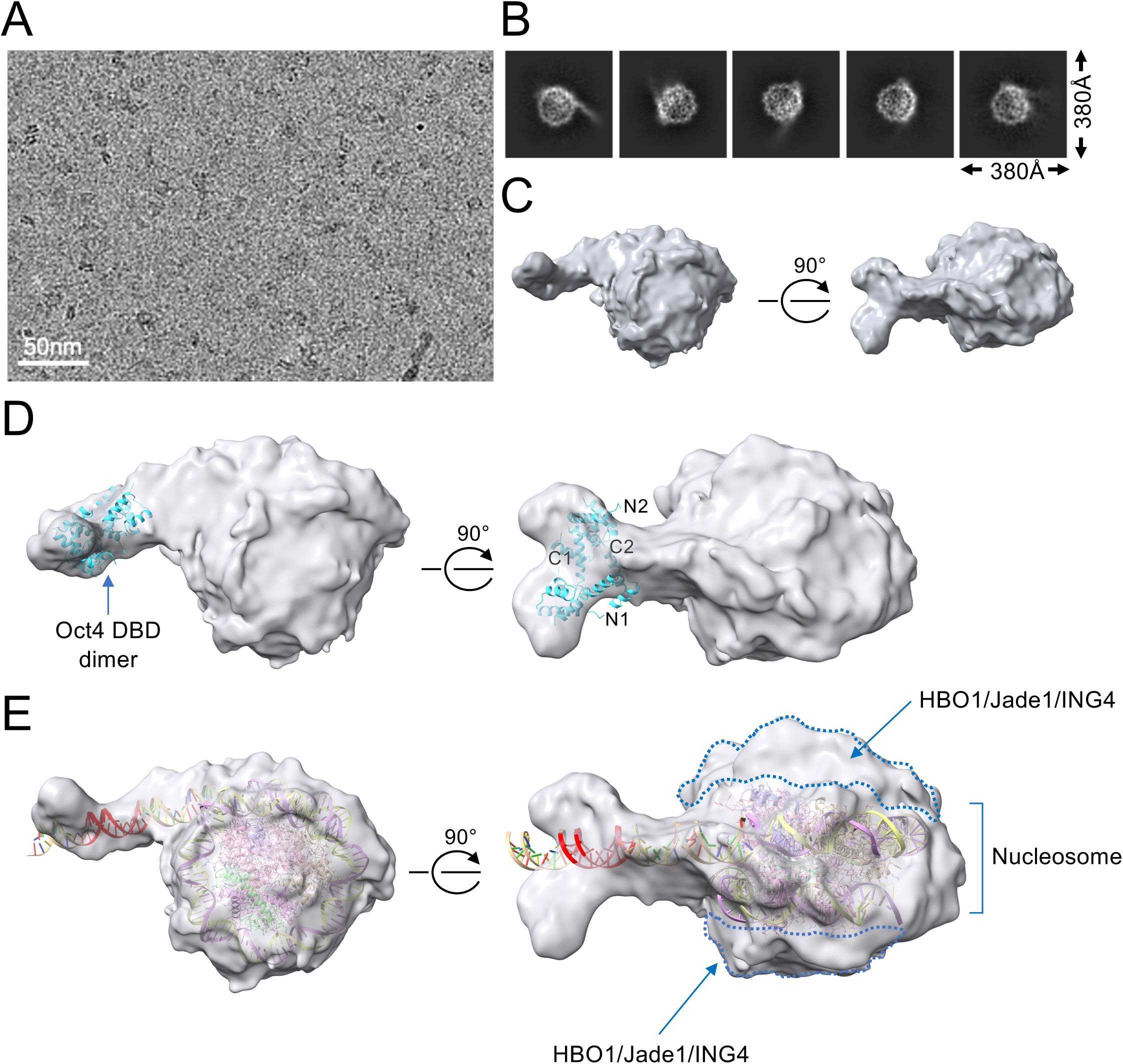
Structural analysis of Oct4 bound to a MORE complexed with HBO1/Jade1 HAT and nucleosome. *A*, Representative cryoEM micrograph of particles generated using HBO1, Jade1, ING4, hEaf6, Oct4 and MORE-containing nucleosomes. *B*, Selected 2D classes are shown. *D*, 3D-reconstructed and refined volume for the HBO1/Jade1/ING4/hEaf6/Oct4/nucleosome complex is shown in two projections rotated 90°. *E*, Two Oct4 DNA binding domain molecules (POUS and POUH, PDB ID: 7XRC) were fitted in the extended volume. Positions of the two DBD N- and C-termini are highlighted. *F*, Partial structure of the nucleosome (PDB ID: 6TEM) fitted within the volume. Part of the DNA was opened and fitted into the extended volume where the predicted location of the MORE is shown in red. Left and right panels were rotated 90°.

## Discussion

The ability of transcription factors bound to DNA in different configurations to interact with distinct cofactors is incompletely understood. Nevertheless, different cofactor specificities can be predicted because proteins associated with DNA in different configurations present different chemical docking surfaces. For example, the Oct4 paralog Oct1 can associate with its B and T cell-specific coactivator OCA-B (OBF-1, Bob.1) when bound to a simple octamer or a PORE but not when bound to a MORE due to occlusion of the docking site in the MORE configuration (10). It has been speculated that POU proteins may provide examples due to their ability to rearrange their DNA binding domains based on the orientation and spacing of their cognate DNA half-sites (32). As a second example, FoxP3 binds to DNA as a dimer in a head-to-head conformation, forming a hydrophobic pocket for Runx1 binding in normal regulatory T cells and leading to immune tolerance. However, some human immunodysregulation polyendocrinopathy enteropathy X-linked (IPEX) patients harbor a *FOXP3* mutation that encodes a protein that binds to DNA in a “forkhead-swapped” configuration in which the DNA binding domains bind DNA in a different configuration eliminating the Runx1 binding pocket (33). Here, we show that the pluripotency master regulator Oct4, bound to DNA in different monomeric and dimeric configurations, selectively associates with distinct cofactors. One of these, the Jade1 component of the HBO1 complex, associates with Oct4 bound as a homodimer bound to DNA sites known as MOREs (Fig. S7).

Prior studies have investigated potential Oct4 interactioning partners (19–21), however, these studies did not specifically analyze interactions with DNA-bound Oct4. Within the genome, Oct4 sites are known to frequently comprise different octamer half-sites with variable numbers, spacing and orientation (34, 35). Oct4 sites are also frequently associated with binding sites for other transcription factors. For example, in pluripotent stem cells a single molecule of Oct4 binds to an Oct:Sox element approximately 2 kb upstream of the *Pou5f1* gene as a heterodimer with Sox2 (36), while four Oct4 molecules are capable of binding upstream of *Polr2a* at two consecutive MOREs (16). We used canonical octamer element or MORE DNA to enrich Oct4 and co-bound proteins from mouse ESC extracts. We opted to use ESCs deficient in the Oct4 paralog Oct1 rather than wild-type ESCs (17, 18) to avoid the identification of confounding proteins uniquely bound by this protein. Analyzing the bound proteins using mass spectroscopy, we identified several proteins consistent with prior reports, including Sox2 and Mta1 (19, 20). Proteins associated with Oct4 bound to canonical octamer sites tended to be associated with transcriptional repression, e.g. Macroh2a1 (37, 38), Ctbp2 (39), Mta1 (40), Kdm2b (41). In contrast, many MORE-associated proteins are associated with transcription activation, e.g. H3.3 (37, 38) and Jade1. This may reflect that fact that Oct4 operating through canonical octamer DNA has been associated with both activation and repression (40, 42), while MORE-containing genes such as *Polr2a*, *Ahcy*, *Rras*, *Ell* and *Taf12* tend to be broadly and strongly expressed. Reporter assays have shown that dimeric Oct4 activates gene expression far more strongly than monomeric Oct4 (11). Prior studies of Oct4 interactors also identified HBO1 (KAT7, MYST2), the catalytic subunit of the HBO1 complex (19).

We propose that Oct4 binding to a MORE orients Oct4 molecules to relieve occlusion of Jade1 by the Oct4 N-termini, whereas Oct4 binding to a canonical site allows for the recruitment of other coregulators, via the N-terminus, to regulate gene expression at pluripotency-associated targets, developmentally poised targets, and other targets lacking MOREs (Fig. S7). Jade1 is a PHD finger-containing scaffolding protein that recognizes histone H3 and helps localize HBO1-contaning HAT complexes to chromatin (24). In human ESCs, HBO1 deficiency is associated with loss of pluripotency, spontaneous neural differentiation, and a defect in mesendoderm specification following TGFβ treatment (43). A role for HBO1 has also been identified in the maintenance of leukemic stem cells (44). We confirmed the Oct4-Jade1 interaction using co-IP in both Oct1 deficient and wild-type ESCs. Because the Oct4 N-terminus has been associated with activation (26), we tested whether the N-terminus was important for this interaction. Oct4 contains a nuclear localization sequence in its homeodomain (27), preserving nuclear localization in the N-terminally truncated Oct4 construct.

Surprisingly, Oct4 lacking its N-terminus showed superior interactions with Jade1. Overexpressed intact Oct4 but not Oct4-ΔN was also sufficient to decrease the protein levels of co-expressed Jade1, and even the Oct4 N-terminus expressed separately was sufficient to achieve this effect. We do not yet know the mechanism underlying this phenomenon. The Oct4 N-terminus is a site of post-translational modification (45, 46), potentially adding another layer of regulation.

HBO1 ChIP-seq studies using Oct1 deficient mouse ESCs identify ∼12,000 binding events and reveal a preference for promoter binding, similar to findings using HBO1 ChIP-seq in HeLa cells (25). Intersecting these peaks with known Oct4 binding sites identifies ∼1,700 co-binding events preferentially enriched for MOREs, which are otherwise rare in the genome compared to canonical octamer elements. In ESCs, HBO1 preferentially associates with DNA adjacent to MOREs at target genes.

In a reconstituted system, purified recombinant Oct4 bound to MOREs adjacent to nucleosomes catalyzes acetylation of H3K9 but not other lysines by HBO1/Jade1 complexes. Lysines in H4 for example were efficiently acetylated by reconstituted HBO1/Jade1-containing complexes, however only histone H3K9 showed augmented acetylation in the presence of Oct4 and MORE DNA. Moreover, conditions that augment Oct4 binding also augment H3K9 acetylation. Prior work showed that MORE binding by the Oct4 paralog Oct1 is inducible through a specific serine residue that when phosphorylated stabilizes a MORE-bound DNA binding configuration, and that a phospho-mimetic mutation augments MORE binding (16). We found that the analogous mutation in Oct4 (Oct4^S229D^) also preferentially bound to MORE DNA. The increase in Oct4 binding was associated with increased HBO1-mediated H3K9 acetylation. The lack of augmentation of H4 acetylation is surprising, however the Oct4 N-terminus is known to bind to the H4 tail (15) and may occlude additional acetylation via this mechanism. Oct4 and its paralog Oct1 are both known to recruit cofactors that act on H3K9 by removing K9me2 marks (47)(48).

Using the same recombinant proteins used in the HAT assay, we determined the cryoEM structure of an Oct4:nucleosome:HBO1 complex at ∼7 Ä resolution, revealing that the complex surrounds both sides of the nucleosome to form a more spherical structure, and revealing that the DNA partially unwinds from the nucleosome core to generate an extension containing the MORE and Oct4 homodimer. Oct4 nucleosome binding is known to be able to alter nucleosomal structure, loosening the DNA winding and allowing for the binding of additional transcription factors, including additional Oct4 molecules (15). Although Oct4 bound to a MORE can catalyze HBO1 complex acetylation, no density was observed adjacent to DNA-bound Oct4. This may be due to flexibility in the Oct4 contact with components of the complex, or the possibility that other structures besides the one refined and described here contain stable Oct4 contacts. Oct4 binding to DNA, particularly through multiple binding sites, is known to induce changes in nucleosome structure, including repositioning of DNA and altering the location of the H3 and H4 tails (15). Cumulatively, these results highlight Oct4 as a prime example of a spatioselective transcription factor that can interact with difference coregulatory proteins when bound to DNA in different configurations.

## Experimental procedures

### Cell culture and conditions

ESCs were cultured on mitomycin C-arrested feeder mouse embryonic fibroblasts (MEFs) in ES medium as described in (47) in the presence supplemented LIF and 2i conditions (3 µM CHIR99021 and 1 µM PD0325901, Selleck Chemicals) at 37°C. One passage prior to experiments, the feeders were depleted by culturing on gelatin coated plates overnight. 3T3 cells were cultured with DMEM medium with 10% FBS, 1.2% Pen Strep (ThermoFisher) and 1x GlutaMAX supplement (ThermoFisher). Cells were transiently transfected using Lipofectamine 3000 (ThermoFisher #L3000-015) according to the manufacturer instructions. Expi293F cells were grown in Expi293 expression media (ThermoFisher #A1435101) in shaker flasks at 37°C in a humidified incubator supplied with 8% CO2 equipped with a shaker set to 125 rpm. Cells were transfected using the Expi293 Expression System (ThermoFisher #A14635) according to the manufacturer instructions. For transfection, Expi293F cells were seeded in 30 mL of Expi293 expression media at a density of 1.25×10^6^ cells/mL in 125 mL flasks. After 24 hr, cells were transfected with wild-type Oct4 expression plasmids (49), Oct4S^229D^ mutant plasmids or Jade1L (30) using an ExpiFectamine 293 Transfection Kit (ThermoFisher) following the manufacturer’s protocol.

### Immobilization of dsDNA on FG beads

Double-stranded DNA (dsDNA) was covalently fixed to FG magnetic beads (Tamagawa-Seiki) using the manufacturer protocol. Briefly, 150 µg of sense and antisense 5’ phosphorylated single-stranded DNA (WT octamer sense: 5’-ACGTATGCAAATAGTGGGGG-3’, WT octamer antisense: 5’-ACTATTTGCATACGTCCCCC-3’; mutant octamer sense: 5’-ACGTA<u>AA</u>C<u>T</u>A<u>C</u>TAGTGGGGG-3’, mutant octamer antisense: 5’-ACTAGTAGTTTACGTCCCCC-3’; WT MORE sense: 5’-CATGCATATGCATGCGGGGG-3’, WT MORE antisense: 5’-GCATGCATATCGATGCCCCC-3’; and mutant MORE sense: 5’-CACACTGTCGTGTGCGGGGG-3’, mutant MORE antisense: 5’-GCACACGACAGTGTGCCCCC-3’ where underlined text indicates the mutation) dissolved in ultrapure water at 1 µg/µL were annealed at 98°C for 10 min and gradually cooled to the room temperature. The annealed dsDNAs were ligated using 700 units of T4 DNA ligase (New England Biolabs) on a rocker platform at 37°C overnight. Ligated DNA was size-selected using a NICK column (GE Healthcare, 200bp-400bp) and eluted in 400 mL ultrapure water. 400 µL DNA (approximately 250 µg/mL) were immobilized to 10 mg of ultrapure water-washed FG beads at 50°C for 24 hr. The beads were then washed with 2.5 M KCl twice, ultrapure water twice and TES buffer (10 mM Tris-HCl pH 8.0, 0.3 M KCl, 1 mM EDTA and 0.02% NaN3) three times. Immobilized beads were dispersed in 400 mL of TES buffer and stored at 4°C. Beads contained at least 2 µg immobilized DNA per mg.

### Oct4 affinity purification

ESCs deficient for the transcription factor Oct1 (17, 18) were pelleted and resuspended in NP-40 lysis buffer (150 mM NaCl, 1% NP-40, 50 mM Tris-Cl pH 8.0) containing cOmplete protease inhibitor cocktail (Roche) and PhosSTOP phosphatase inhibitor cocktail (Roche). Cells were lysed by rotating in the buffer and clarified by centrifugation at 13,000×*g* at 4°C. Protein concentrations were normalized using Bradford assays (Bio-Rad) and diluted (>5-fold) to 1 mg/mL using 100 mM KCl buffer (100 mM KCl, 10% glycerol (v/v), 20 mM HEPES pH 7.9, 2 mM MgCl2, 0.2 mM CaCl2, 0.2 mM EDTA, 0.1% NP-40, 1 mM dithiothreitol) with protease and phosphatase inhibitors. DNA-immobilized FG beads were washed with 100 mM KCl buffer, and 0.5 mg of the beads were added to the diluted cell lysates and rotated at 4°C for 4 hr. The beads were sequentially washed 3 times with 100, 200 and 300 mM KCl buffer. Bound proteins were eluted with 300 mM KCl buffer containing 30 µM of the corresponding free DNA oligomers for 1 hr at 4°C on a rotating platform. For experiments testing Oct4 lacking its N-terminus, lysates contained 20 μg/mL MG-132 (Sigma) and post-binding to MORE DNA-conjugated FG beads were washed three times with 100 mM KCl buffer.

### Western blotting

Jade1L, Jade1S, HBO1, ING4 and hEaf6 proteins were detected using antibodies from Novus (NBP1-83085), Proteintech (15032-1-AP), Abcam (ab124993), Abcam (ab108621) and Novus (NBP1-91523) respectively. Recombinant Oct4 and Oct4 S229D were detected using an antibody from Santa Cruz (sc-5279). Twin-strep-tagged proteins were detected using HRP conjugated anti-StrepII antibody (Genscript). Acetylated Histone H4 was detected by Abcam H4 antibody cat no ab177790. Histone H3 was detected using an antibody from Cell Signaling Technology (CST, 4499T). H4K5, H4K8, H4K12, H4K16, H3K9, K3K14, H3K18 and H3K27 acetylation were detected using monoclonal antibodies from CST (8647T, 2594T, 13944T, 13534, 96495, 7627T, 13998T and 8173T respectively).

### Mass spectrometry and analysis

Proteins eluted from DNA-bound FG beads above were resolved by 10% SDS-PAGE approximately 2 cm unto the running gel and stained with Colloidal Blue (Thermo Scientific). The gel slabs were excised and destained until they were clear. Proteins in the gel slabs were reduced with 25 mM DTT at 37°C for 30 min, followed by alkylation with 50 mM IAA for 45 min at room temperature. Proteins were then digested with 1.0 µg of trypsin overnight at 37°C. The peptides were extracted from the gel, dried completely, desalted and suspended in 20 µL of 0.1% formic acid for LC-MS/MS analysis. Reverse-phase nano-LC/MS/MS was performed on an UltiMate 3000 RSLCnano system (Dionex) coupled to a Thermo Scientific Q Exactive-HF mass spectrometer equipped with a nanoelectrospray source. Peptides were diluted to 0.2 µg of sample in 0.1% formic acid in water. 5 µL of the samples were injected onto the liquid chromatograph. A 50-cm long/100 µm inner diameter reverse-phase BEH C18 3.0 µm nanocolumn (Thermo Scientific) was employed for chromatographic separation. A gradient of buffers (Buffer A: 0.1 % formic acid in water; Buffer B: 0.1% formic acid in acetonitrile) at a flow rate of 200 µL/min at 60°C was used. The LC run lasted for 83 min with a starting concentration of 5% buffer B increasing to 55% over the initial 53 min and a further increase in concentration to 95% over 63 min. MS/MS data was acquired using an auto-MS/MS method selecting the 20 most abundant precursor ions for fragmentation. The mass-to-charge range was set to 350–1800. For these samples, the Uniprot database was searched with the *Mus musculus* taxonomy selected. An allowance was made for 2 missed cleavages following trypsin/Lys-C digestion. The fixed modification was cysteine carbamidomethylation. The variable modifications were methionine oxidation, and N-terminal deamidation. The search was performed with a mass tolerance of 15 ppm for precursor ions and a mass tolerance of 10 ppm for fragment ions. Protein enrichment was analyzed by MASCOT using N=3 biological replicates. The raw data were processed by MetaboAnalyst (50), and the missing values were estimated by KNN. Raw data were processed with log transformation and Pareto scaling. The threshold was set as 2-fold enrichment and *p*<0.05.

### Oct4 N-terminal deletion

Oct4 fused with C-terminal twin-strep and FLAG tags cloned into the pACE-MAM2 vector (Geneva Biotech) for mammalian cell expression was described previously (49). The N-terminal Oct4 truncation lacking the first 130 amino acids was generated by PCR amplification with primers 5‘-ctcaatcCATGGACATGAAAGCCCTGCAG-3‘ and 5‘-GAATGCATCAGCTGGTTTGAATGCATGGG-3‘ where the lower-case letters indicate divergence from the parent vector. The wild-type and N-terminal-truncated Oct4 PCR fragments were digested with *Nco*I and *Pvu*II and ligated back into the pACE-MAM2 vector digested with *Nco*I and *Pvu*II to generate the truncated Oct4 vector. All constructs were sequenced to confirm veracity.

### Co-IP

ESCs and 3T3 cells transfected with the Oct4 or Oct4ΔN plasmids described above were lysed with RIPA buffer (150mM NaCl, 50mM Tris-HCl pH 8.0, 1% NP-40, 0.5% sodium deoxycholate, 0.1% SDS) containing cOmplete protease inhibitor cocktail (Roche) by rotating at 4°C for 30 min.

Lysates were clarified by centrifugation (13,000×*g*) for 10 min. 2 µg of primary antibodies were added to the lysates, which were rotated at 4°C overnight. Antibodies used were as follows: anti-Oct4 ×Santa Cruz sc-5279), anti-Jade1 (Proteintech 15032-1-AP), anti-FLAG M2 (Sigma-Aldrich F1084). Isotype control IgG was purchased from Jackson ImmunoResearch. 20 µL RIPA buffer-washed Protein G Dynabeads (Thermo Scientific 10004D) were added to the cell lysis-antibody mix, which were rotated at 4°C for an additional 5 hr. Protein-bound Dynabeads were then washed with 500 µl RIPA buffer for 4 times, and proteins were eluted with 50 µL 2x SDS loading buffer (4% SDS, 20% glycerol, 0.2% bromophenol blue, and 200 mM DTT) by heating at 98°C for 5 min. For the separate expression of the recombinant Oct4 N-terminus, amino acids 1-130 were synthesized as an g-block (IDT). The Oct4 expression vector was digested with *Nco*I and *Nsi*I to remove the full-length Oct4 cassette, and NEBuilder HiFi DNA Assembly (New England Biolabs) was used to insert the Oct4 N-terminus g-block into the digested vector.

### ChIP-seq

ChIP-seq was performed as previously described (18). Briefly, Oct1 deficient ESCs were crosslinked with 1% formaldehyde, quenched with 2.5 M glycine, and crude nuclei were extracted using Farnham lysis buffer (5 mM PIPES pH 8.0, 85 mM KCl, 0.5% NP-40), and further lysed with RIPA lysis buffer. Chromatin was sheared on ice using EpiShear probe sonicator (Active Motif) set to amplitude=40, 30 sec on/30 sec off for 5 min. 4 μg of rabbit anti-HBO1 antibody (Cell Signaling #58418S) was applied to the sheared chromatin and the mixture incubated at 4°C overnight. 1.5 mg of protein G Dynabeads (Thermo Scientific) were applied to samples, which were rotated at 4°C for 5-6 hr to allow binding, and the beads were collected and washed with Low Salt buffer (20 mM Tris-Cl pH 8.0, 150 mM NaCl, 2 mM EDTA, 0.1% SDS, 1% Triton X-100), High Salt buffer (identical but with 500 mM NaCl), Low Salt buffer, LiCl buffer, and TE buffer (Tris-EDTA pH 8.0 plus 1 mM EDTA) for 5 washes in total, each with 10 min rotating at 4°C. Chromatin was eluted from the beads using IP Elution buffer (1% SDS and 0.1M NaHCO3) and incubated at 65°C overnight to reverse the formaldehyde crosslinks. Before IP, 2% Input was reserved from the samples. DNA was purified from both IP samples and input using phenol-chloroform-isoamyl alcohol extraction followed by QIAquick PCR purification kit (Qiagen). N=2 biological repeats were performed per condition. ChIP materials were sequenced using a NovaSeq instrument (Illumina).

### ChIP-seq analysis

Between 16 and 25 million paired-end reads were aligned to the mouse Mm10 reference genome using Novoalign v4.03.01. HBO1 and Oct4 ChIP-seq data was analyzed using Macs2 (51) with parameters of q 0.005, max-gap 100 and q 0.005, max-gap 100, min-length 200 respectively. HBO1 and Oct4 ChIP peak summits within 100 bp were defined as overlapping peaks. Oct4 peaks and overlapping peaks were analyzed with MEME-chip CentriMo (52) for motif enrichment. The raw and processed ChIP-seq data have been deposited with the gene expression omnibus (GEO, series record GSE279191).

### Recombinant Oct4 and Jade1 purification

Wild-type Oct4 expression plasmids (49) were transfected into Expi293F cells (Thermo Scientific) using the ExpiFectamine 293 Transfection Kit (Thermo Scientific) following the manufacturer instructions. 48 hr post-transfection, cells were harvested and washed with ice-cold PBS. Cells were resuspended in Buffer A (20 mM HEPES pH 8.0, 1.5 mM MgCl2, 10 mM KCl, 0.25% NP-40, 0.5 mM dithiothreitol) containing protease inhibitor cocktail (Roche), incubated for 10 min on ice, and homogenized using a Dounce homogenizer (Wheaton). The homogenate was centrifuged at 4°C for 5 min at 3800×*g* to pellet nuclei, which were resuspended in Buffer C (20 mM HEPES pH 8.0, 25% glycerol, 1.5 mM MgCl2, 420 mM KCl, 0.25% NP-40, 0.2 mM EDTA, 0.5 mM dithiothreitol) with protease inhibitors and again Dounce homogenized. Nuclei were extracted for 30 min at 4°C with end-to-end rotation, then centrifuged at 4°C for 30 min at 20,000×*g*. The nuclear extract was then immunoprecipitated with anti-FLAG (M2)-agarose beads (Sigma). Bead-bound proteins were eluted with 200 ng/µL 3’FLAG peptide (Sigma) by incubating at 4°C for 30 min. To generate an Oct4 point mutation corresponding to Oct1 S385D (16), Oct4 S229 was mutated by overlap extension PCR using as middle primers 5’-GAGAAAGCGAACTGATATTGAGAACCG-3’, 5’-CGGTTCTCAATATCAGTTCGCTTTCTC-3’, and as end primers 5’-GGATCTCGAGCCATGGCTGG-3’, 5’-GAATTAGGTACCTTATTTGTCATCATCG-3’. For Jade1 purification, 60 µg of full-length Jade1 expression plasmid (30) (a gift of Dr. Jacques Côté) was used and purified similarly. All proteins were pure and monodisperse as verified using SDS-PAGE/Coomassie blue staining and size exclusion chromatography.

### Baculovirus expression for cryoEM

The pFastBac-1 vector was digested with *BamHI* and *EcoRI* and ligated with annealed oligos 5’-GATCCATGTCGCATCACCATCACCATCACAGGCTCGAGGCCATGGTG-3’ and 5’-AATTCACCATGGCCTCGAGCCTGTGATGGTGATGGTGATGCGACATG-3’ to incorporate a 6-His tag as well as *Xho*I and *Nco*1 sites. A single base frameshifted version of pFastBac-1 was also generated using the oligos 5’-GATCCATGTCGCATCACCATCACCATCACGAGGCTCGAGGCCATGGTG-3’ and 5’-AATTCACCATGGCCTCGAGCCTCGTGATGGTGATGGTGATGCGACATG-3’. cDNAs encoding wild-type or S229D Oct4 with C-terminal twin-strep and FLAG-tags were then subcloned into the modified pFastBac-1 vector between the *Nco*I and *Kpn*I sites. The *Ing4* cDNA was amplified using PCR from a plasmid template (Addgene #13289) using 5’-AGGCTCGAGATGGCTGCGGGGATG-3’ and 5’-AGAATGCATTTTCTTCTTCCGTTCTTGG-3’ primers and was cloned into the *Xho*I and *Nsi*I sites using the Oct4/pFastBac-1 vector by first excising the Oct4 cDNA using *Xho*I and *Nsi*I, preserving the N-terminal His-tag and C-terminal twin-strep tags. HBO1 and hEaf6 gene blocks containing C-terminal twin-strep-tag were synthesized from IDT and cloned into the modified single-base frameshifted pFastBac1 vector using the *Xho*I and *Kpn*I sites. Full-length Jade1L was cloned in the backbone of the Oct4 expression plasmids (49) using the *Xho*I and *Nsi*I sites to replace Oct4, then Jade1L with a C-terminal twin-strep tag was amplified by PCR from the Jade1L mammalian cell expression plasmid using the primers sets 5’-GATCTCGAGCCATGAAACGAGGTCGCCTTCC-3’ and 5’-TATGCATGCTTATTTGTCATCATCG-3’ and cloned into the *Xho*I and *SphI* sites of the modified single-base frameshifted pFastBac-1 vector. Constructs were transposed into bacmids using DH10Bac competent cells (Thermo Scientific #10361012). Transposed recombinant bacmids were purified using a PureLink plasmid isolation kit (Thermo Scientific #K210004). Transposition was confirmed by PCR. Recombinant bacmids were transfected into Sf9 cells to generate recombinant virus and amplified for large scale protein purification by growth to 3-4×10^6^ cells/mL in Sf-900 II SFM media (ThermoFisher #10902088) in a humidified 28°C incubator shaking at 125 rpm. Cells were collected, added to fresh media at 2×10^6^ cells/mL, and infected with recombinant baculovirus. Cells were grown for 48 hr, harvested by centrifugation and lysed in lysis buffer (20 mM HEPES pH 8.0, 300 mM NaCl, 5% glycerol, 0.5mM DTT) containing protease inhibitor cocktail (Roche) using an Active Motif probe sonicator. The lysate was centrifuged at 22700×*g* for 30 min. The supernatant was loaded onto a 1 mL prepacked Strep-tactin XT column (Cytiva #29401317), washed with 10 column volumes of wash buffer (20 mM HEPES pH 8.0, 300 mM NaCl, 5% glycerol, 0.5 mM DTT), and eluted with wash buffer containing 50 mM biotin (Sigma). Purity was confirmed by 12% SDS-PAGE and Coomassie staining.

### Nucleosome preparation

153 bp DNA constructs containing either MORE (ATCCTGGAGAAATGCATATGCATGGCCGCTCAATTGGTCGTAGACAGCTCTAGCACCGCTTAAACGCACGTACGCGCTGTCCCCCGCGTTTTAACCGCCAAGGGGATTACTCCCTAGTCTCCAGGCACGTGTCAGATATATACATCCTGTGAT) or octamer sequence (ATCCTGGAGAATCCCGGATGCAAATCCGCTCAATTGGTCGTAGACAGCTCTAGCACCGCTTAAACGCACGTACGCGCTGTCCCCCGCGTTTTAACCGCCAAGGGGATTACTCCCTAGTCTCCAGGCACGTGTCAGATATATACATCCTGTGAT) were designed based on the octamer binding site as previously described (14). 16 copies of the DNA were cloned into the pUC19 vector using published methods (53). The plasmid containing 16 copy of desired DNA fragment was prepared by large scale plasmid purification followed by *Eco*RV-HF digestion. The final fragment was purified by PEG8000 precipitation (54). Recombinant human histones H2A, H2B, H3.3 and H4 were expressed from transfected plasmids (55) (a gift of Dr. Brad Cairns) and individually purified from Rosetta2 DE3 cells (Sigma-Aldrich #71403) and histone octamer was reconstituted and purified using published protocols (56). Reconstituted octamers were purified by size exclusion chromatography using a Superdex 200 Increase 10/300 GL column (Cytiva #28990944). Nucleosomes were reconstituted using a 1:1.1 molar ratio of histone octamer to DNA and purified by ion exchange chromatography following the protocol described (57), then dialyzed into 20 mM HEPES, pH 7.4 with 50 mM KCl.

### HAT assay

HAT assays were performed in 20 µL reactions containing 225 nM purified nucleosomes, 200 ng purified Jade1L/HBO1 complex, and 200 µM acetyl-CoA (Sigma #A2181) ±250ng of Oct4 in 20 mM Tris pH 7.4, 40 mM KCl and 5mM MgCl2. Reactions were incubated at 37°C for 30 min. Reactions were quenched by the addition of 5.5 mL of 5X SDS-PAGE sample buffer. Samples were boiled at 98°C for 10 min and analyzed by Western blot using specific antibodies. The extent of acetylation was quantified by ImageJ software (National Institutes of Health).

### Jade1 knockdown

SMART pool ON-TARGETplus mouse Jade1 siRNAs were purchased from Dharmacon (#269424). Cultured ESCs were first depleted of feeders by plating on gelatin for 30 min. Cells were then removed to new gelatin-coated plates, and transfected 12 hr later with Lipofectamine 3000 (ThermoFisher). After an additional 48 hr, cells were processed for immunoblot or ChIP.

### ChIP-qPCR

ChIP assays were conducted identically to the ChIP-seq outlined above and antibodies against H3K9Ac (Abcam ab4441). qPCR conditions and primers for *Polr2a*, *Pou5f1* and 28S were identical to those used in (18).

### CryoEM and image processing

The HBO1 complex was reconstituted at 1.5 µM final concentration in 20 mM HEPES, pH 7.4, 50 µM KCl and 1 mM DTT using individually purified HBO1, Jade1L, ING4 and hEaf6 from Sf9 cells (ATCC #CRL-1711) and incubated over ice for 15 min. 1.5 µM of purified Oct4^S229D^ was separately combined with an equimolar amount of purified nucleosome core particles and incubated over ice for 15 min. The reconstituted HBO complex was then added to the Oct4/nucleosome complex and incubated over ice for an additional 30 min. The reconstituted complex was then lightly crosslinked using 0.01% (v/v) glutaraldehyde (Sigma) for 30 sec. Excess crosslinker was quenched using Tris-Cl pH 7.5. The final complex was concentrated to ∼0.5 mg/mL using a 0.5ml Amicon ultra-centrifugal filter with a cutoff 10 kD (Sigma Millipore). 3.5 µL of concentrated complex was added to a glow-discharged UltraAufoil Gold Holey Films (R1.2/1.3 300-mesh Au, EMS #Q350AR13A). After blotting the grids for 4 sec at 4°C and 95% relative humidity, the samples was plunged into liquid ethane using a Vitrobot Mark IV (ThermoFisher). CryoEM images were recorded using a Titan Krios 300 kV microscope equipped with a Gutan K3 camera at 80000X magnification for a final pixel size 1.06 Å/pixel with a defocused range -0.8 to -1.8 μm using EPU data acquisition software (ThermoFisher). Images were recorded under low-dose conditions with a total dose of ∼ 60 electrons per Å^2^ over 60 frames. A total of 6200 dose-fractionated image stacks were processed using CryoSPARC 4.0 (58). Images were corrected by Patch Motion Correction, then contrast transfer function (CTF) was estimated by Patch CTF Estimation. The dose-fractionated image stacks were the manually curated to improve the quality of data. Particles were then manually picked, and templates were created for template-based particle picking using box size 380 Å^3^. A total of 2×10^6^ particles were parsed into 2D classes. 100782 particles were selected from the 2D classes for *ab initio* reconstruction, and a heterogeneous refinement was performed using the reconstructed 3D volume. Each group of particles from heterogeneous refinement was taken for non-uniform (NU) refinement. The best resolution obtained (gold-standard Fourier shell correlation, GSFSC) was 7.2 Å with a B factor of 322. The volume map from the best resolution was smoothened using UCSF chimera at a standard deviation of 1.5. The crystal structure of the Oct4 DNA binding domain bound to a MORE (PDB ID: 7XRC) (59) and part of the nucleosome crystal structure (PDB ID: 6TEM) was manually fitted using UCSF chimera. Part of the DNA from the same nucleosome was removed and replaced with B-form DNA of the correct sequence generated separately using UCSF Chimera X. The processed data have been uploaded to the Electron Microscopy Data Bank (EMDB, deposition code EMD-47299).

## Data availability

Mass spectrometry data have been uploaded to the MassIVE database (MSV000096131).

ChIPseq data are available at the Gene Expression Omnibus (GEO) under series record GSE279191). The cryoEM data have been uploaded to the Electron Microscopy Data Bank (EMDB, code EMD-47299.

## Acknowledgements

We thank Jacques Cotê for the gift of the Jade1L expression construct. We thank J. Cox and the Proteomics core facility for assistance with mass spectrometry. We thank B. Dalley and the High-Throughput Sequencing Core facility, and T. Parnell and the Bioinformatics core facility for assistance with ChIP-seq and analysis. We thank D. Belnap, B. Ganser-Pornillos and the Electron Microscopy Core facility for assistance with cryoEM. We thank B. Cairns and members of his laboratory for the gift of recombinant histone plasmid constructs.

## Author contributions

Y.W., A.M., L.L., H.H., M.B.C., D.T.-manuscript writing; Y.W., A.M., Z.S. and M.B.C.-figure generation; Y.W., A.M., L.L., Z.S. and M.B.C.-experiments and data analysis; Y.W., A.M., H.H. and M.B.C.-generation of new reagents for the study; Y.W., A.M. and D.T.-experimental design; D.T.-conception.

## Funding and additional information

This work was supported by a grant from the NIH (1R03TR004568) to DT. We acknowledge indirect support from the Huntsman Cancer Institute’s Cancer Center Support Grant.

## Abbreviations

ChIP-seq: chromatin immunoprecipitation and sequencing
co-IP: co-immunoprecipitation
CryoEM: Cryo-electron microscopy
CTF: contrast transfer function
dsDNA: double-stranded DNA
EMDB: electron microscopy data bank
ESC: embryonic stem cell
GEO: gene expression omnibus
GSFSC: gold-standard Fourier shell correlation
HAT: histone acetyltransferase
IPEX: immunodysregulation polyendocrinopathy enteropathy X-linked
MEF: mouse embryonic fibroblast
MORE: more palindromic octamer related element
NCP: nucleosome core particle
NU: non-uniform
PORE: palindromic octamer related element
POU: Pit-Oct-UNC
POUH: POU homeodomain
POUS: POU-specific domain
TSS: transcription start site.

## Supplemental Figure Legends

**Figure S1. Analysis of mass spectrometry hits from Oct4 affinity purifications and validation of Jade1 binding.** *A*, Venn diagram comparing unique and common identified proteins enriched using affinity purification with octamer compared to MORE DNA. Percentage of protein hits unique to the two conditions are shown. Example proteins from each of the three groups are shown. *B*, The union of all identified proteins enriched using octamer or MORE affinity purification (78 total proteins) was compared with proteins identified from three prior studies (19–21). Proteins previously identified in the different studies are shown. Bold text indicates proteins selectively enriched using MORE DNA in (A). Figure adapted from Ding et al., 2011. *C*, Wild-type ESC lysates were immunoprecipitated with an Oct4 antibody and corresponding mouse IgG as negative control and immunoblotted with Jade1 antibodies.

**Figure S2. Characterization of HBO1 binding events in ESCs.** *A*, Analysis of HBO1 ChIP peak genomic locations based on *Mm10* genome region annotations. *B*, HBO1 ChIP peaks intensities were aligned based on transcription start site (TSS) and plotted based on ±distance from the TSS. *C*, The distance between Oct4 and HBO1 ChIP-seq peak centers was calculated for all common genes (N=2870), and for MORE-containing genes (*Polr2a*, *Zmiz*, two peaks in *Blacap*, and *Rras2*). The average interval is plotted with mean and ±SEM. SEM was used instead of standard deviation because of the highly unequal N values.

**Figure S3. Purification of recombinant components of the HBO1 complex.** *A*, Individual proteins were overexpressed in Expi293F cells and purified using their FLAG tags. The purified proteins were analyzed by 12% SDS-PAGE and Coomassie staining. Arrow indicates the position of the specific protein. *B*, Proteins copurifying with Jade1L were confirmed by Western blot using the corresponding antibodies.

**Figure S4. Recombinant protein purification of Jade1L, HBO1, ING4 and hEaf6.** *A*, Jade1L and HBO1 were independently purified from sf9 cells using their twin-strep tags. Purified proteins were analyzed by 12% SDS-PAGE. Arrows indicate the position of the specific protein in the Coomassie stained gels. *B*, Similar purification of ING4 and hEaf6. Oct4^S229D^ used in cryoEM was overexpressed and purified from Expi293F cells using FLAG tags.

**Figure S5. Design and activity of recombinant Oct4 protein with augmented MORE binding characteristics.** *A*, Sequence alignment of human Oct1 (accession number: AAH52274.1) and Oct4 (accession number: NP_038661.2) is shown. Red rectangle indicates S385 in Oct1, which corresponds to position S229 in Oct4. *B*, HAT assay similar to Fig. 5E except Jade1S was used instead of Jade1L. *C*, Quantification from N=3 experiments using the ratio of acetylated H3K9 to H4 as an internal standard. Error bars denote ±standard deviation.

**Figure S6. Workflow for cyro-EM analysis.** *A*, Schematics for cryoEM data analysis using CryoSPARC. The number of the particles and resolution are shown under each group. *B*, Real space slices are shown for the corresponding NU refinement. *C*, GSFSC resolution plots are shown for the corresponding NU refinement. *D*, viewing direction distributions are shown for the corresponding NU refinement.

**Figure S7. Model for spatioselective recruitment of Jade1 and HBO1 by multimeric Oct4 at nucleosomes to acetylate H3K9.** *A*, At typical binding sides, Oct4 recruits more generic cofactors (X and Y) to regulate transcription either positively or negatively. *B*, At MOREs, dimeric Oct4 associates with Jade1 and the HBO1 HAT complex to acetylate H3K9 at local nucleosomes. Other components of the complex such as Eaf6 and ING4 are also shown.

