## Supplemental figures and legends for "Jade1 and the HBO1 histone acetyltransferase complex are spatial-selective cofactors of the pluripotency transcription factor Oct4"

Supplemental material includes:
Supplemental Figs. S1-S7
Supplemental Tables S1-S3 (provided in separate .xlsx files)

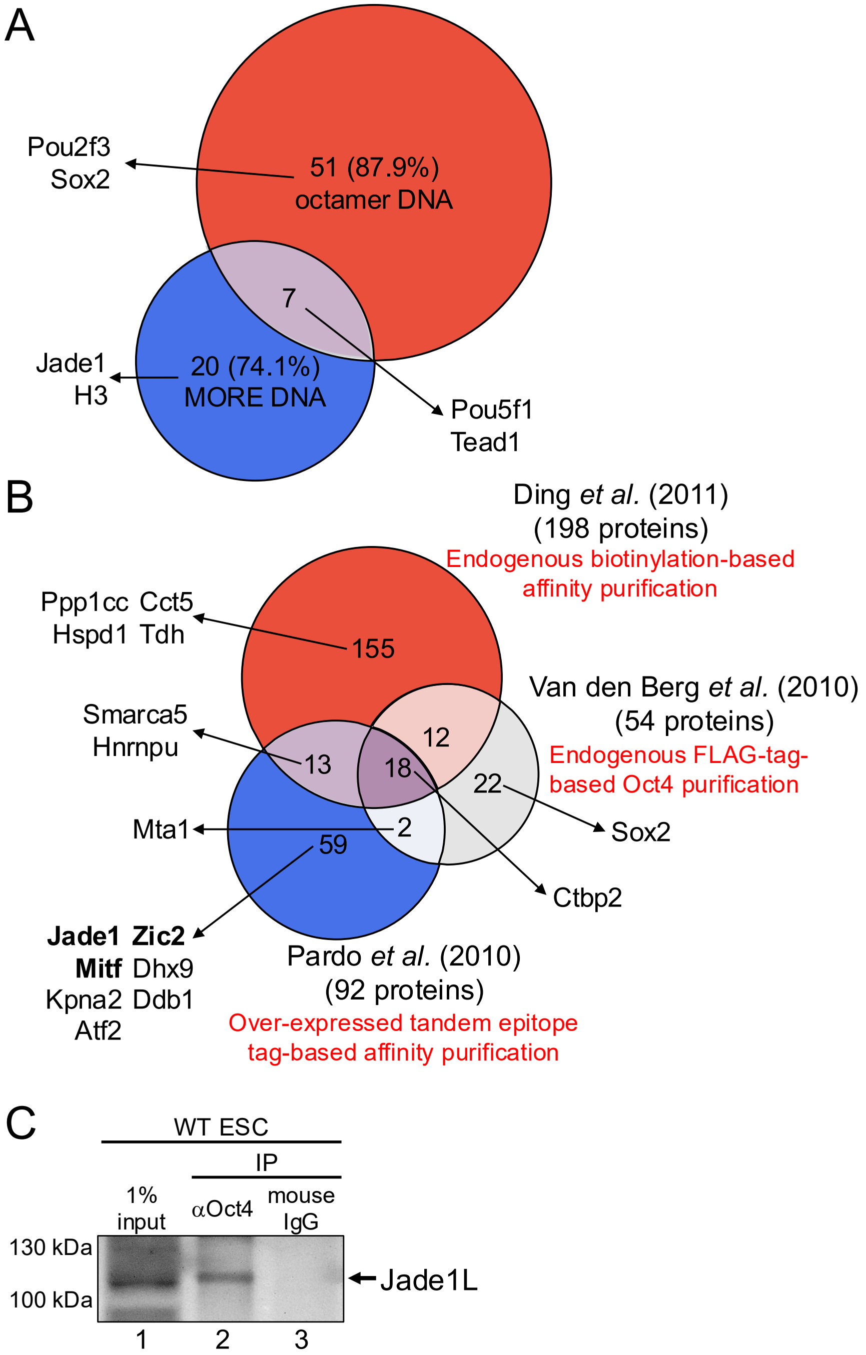


**Supplemental Figure S1. Analysis of mass spectrometry hits from Oct4 affinity purifications and validation of Jade1 binding.** *A*, Venn diagram comparing unique and common identified proteins enriched using affinity purification with octamer compared to MORE DNA. Percentage of protein hits unique to the two conditions are shown. Example proteins from each of the three groups are shown. *B*, The union of all identified proteins enriched using octamer or MORE affinity purification (78 total proteins) was compared with proteins identified from three prior studies (Pardo et al. 2010; van den Berg et al. 2010; Ding et al. 2012). Proteins previously identified in the different studies are shown. Bold text indicates proteins selectively enriched using MORE DNA in (A). Figure adapted from Ding et al., 2011. *C*, Wild-type ESC lysates were immunoprecipitated with an Oct4 antibody and corresponding mouse IgG as negative control and immunoblotted with Jade1 antibodies.

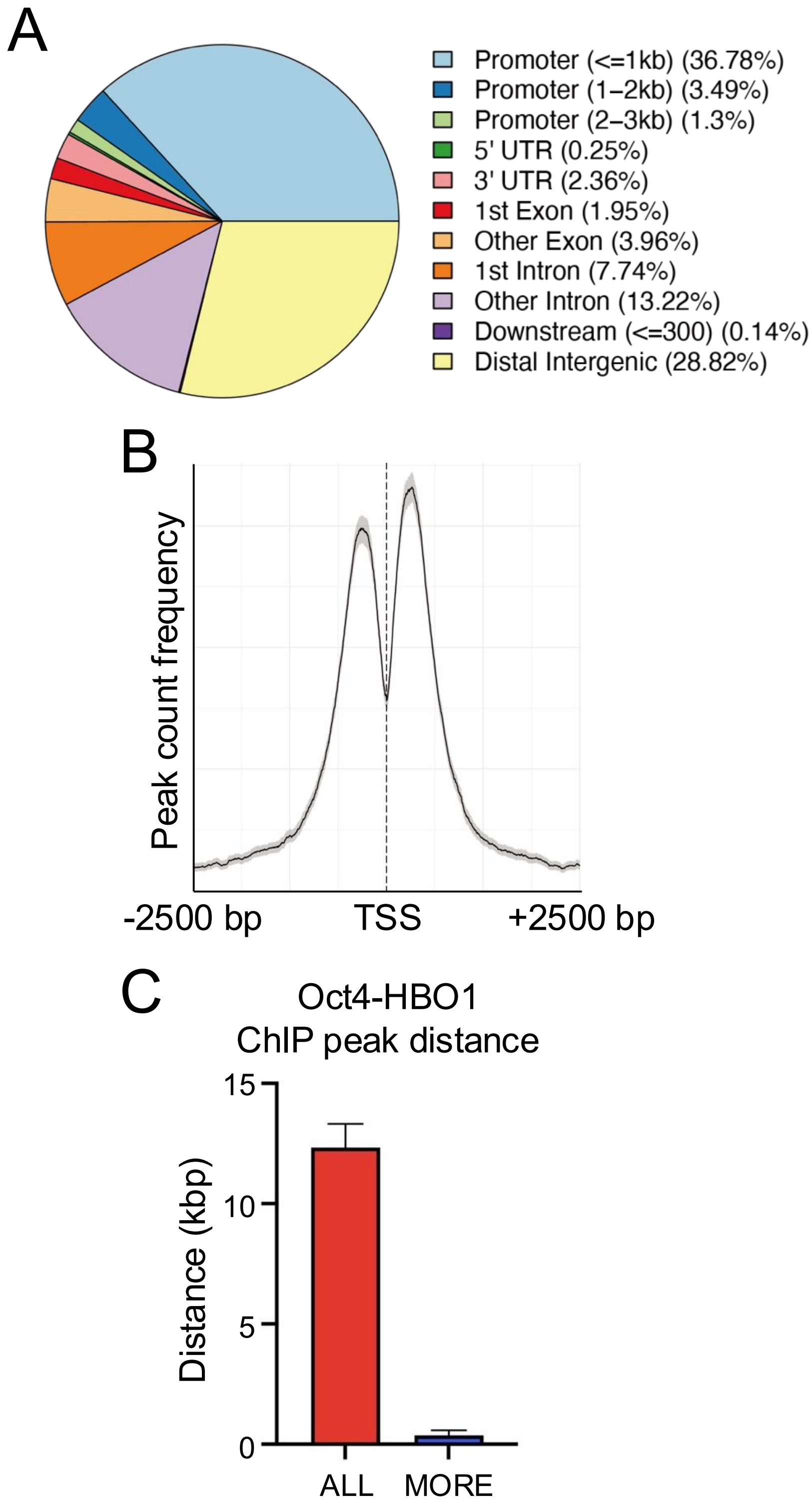


**Supplemental Figure S2. Characterization of HBO1 binding events in ESCs.** *A*, Analysis of HBO1 ChIP peak genomic locations based on *Mm10* genome region annotations. *B*, HBO1 ChIP peaks intensities were aligned based on transcription start site (TSS) and plotted based on ±distance from the TSS. *C*, The distance between Oct4 and HBO1 ChIP-seq peak centers was calculated for all common genes (N=2870), and for MORE-containing genes (*Polr2a*, *Zmiz*, two peaks in *Blacap*, and *Rras2*). The average interval is plotted with mean and ±SEM. SEM was used instead of standard deviation because of the highly unequal N values. SEM was used instead of standard deviation because of the highly unequal N values.

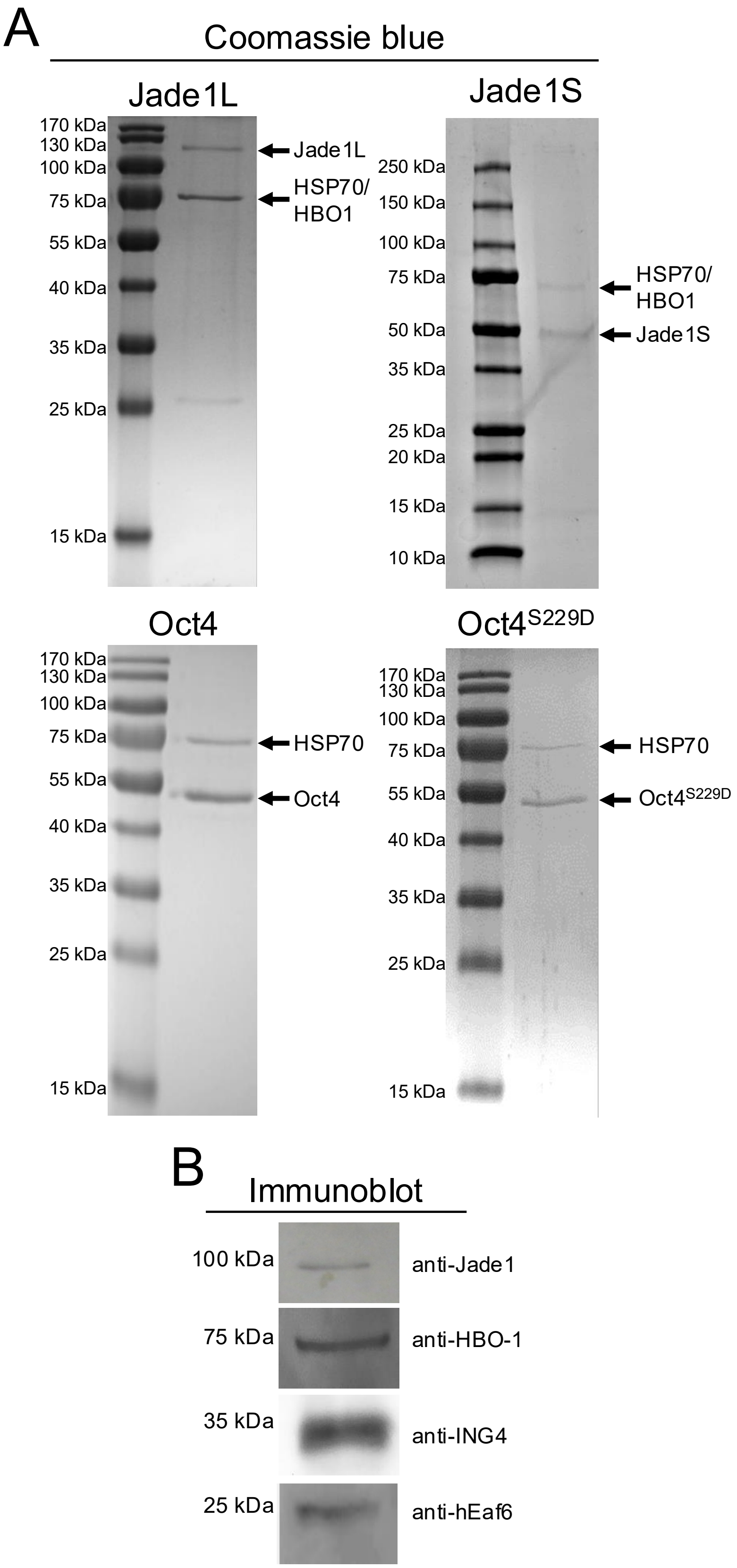


**Figure S3. Purification of recombinant components of the HBO1 complex.** *A*, Individual proteins were overexpressed in Expi293F cells and purified using their FLAG tags. The purified proteins were analyzed by 12% SDS-PAGE and Coomassie staining. Arrow indicates the position of the specific protein. *B*, Proteins copurifying with Jade1L were confirmed by Western blot using the corresponding antibodies.

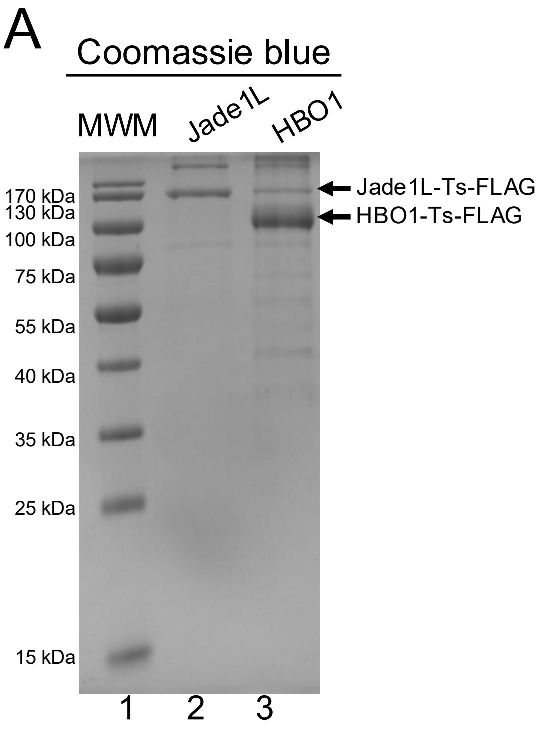


**Figure S4. Recombinant protein purification of Jade1L, HBO1, ING4 and hEaf6.** *A*, Jade1L and HBO1 were independently purified from sf9 cells using their twin-strep tags. Purified proteins were analyzed by 12% SDS-PAGE. Arrows indicate the position of the specific protein in the Coomassie stained gels. *B*, Similar purification of ING4 and hEaf6. Oct4$^{S229D}$ used in cryoEM was overexpressed and purified from Expi293F cells using FLAG tags.

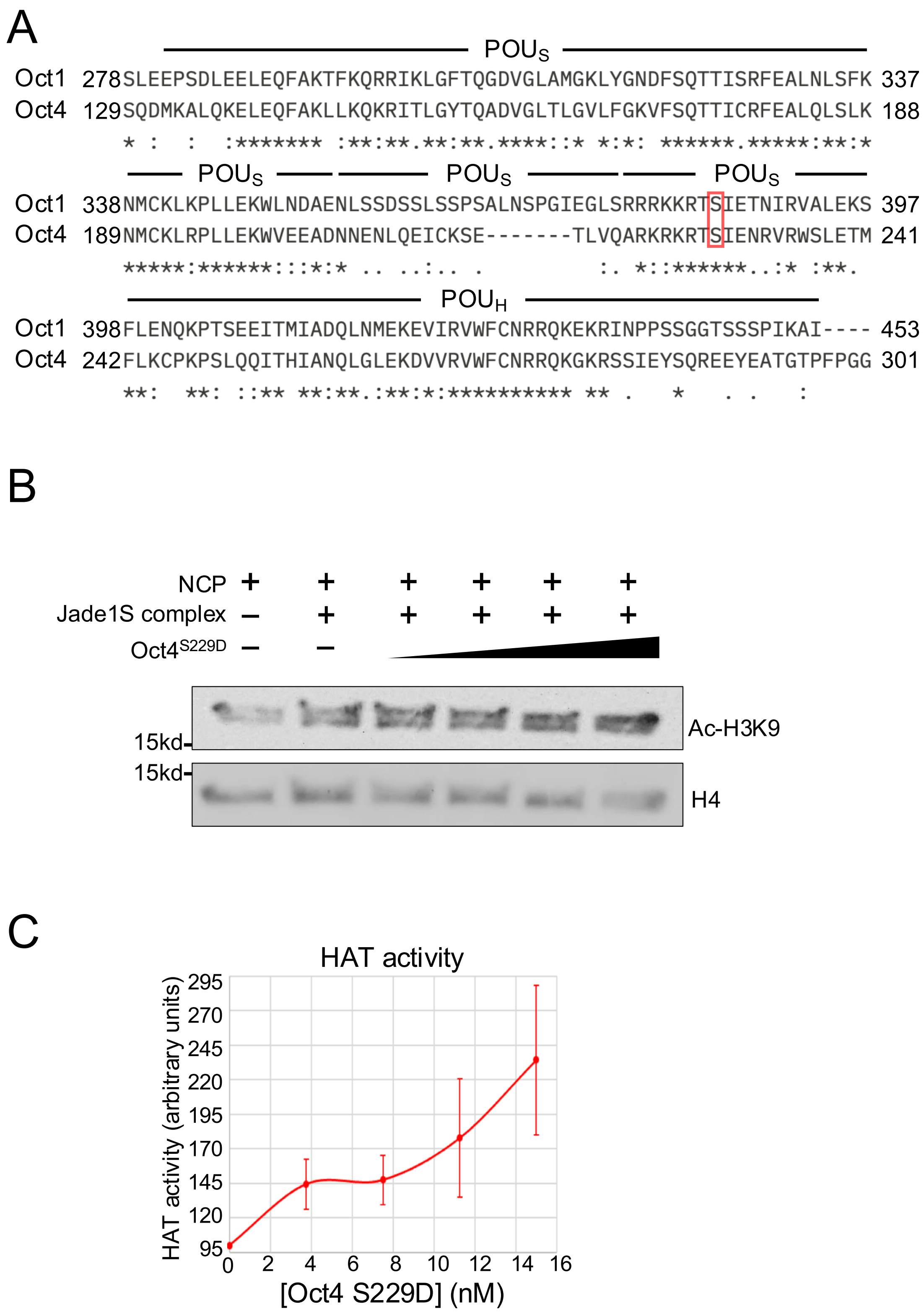


**Figure S5. Design and activity of recombinant Oct4 protein with augmented MORE binding characteristics.** *A*, Sequence alignment of human Oct1 (accession number: AAH52274.1) and Oct4 (accession number: NP_038661.2) is shown. Red rectangle indicates S385 in Oct1, which corresponds to position S229 in Oct4. *B*, HAT assay similar to Fig. 5*E* except Jade1S was used instead of Jade1L. *C*, Quantification from N=3 experiments using the ratio of acetylated H3K9 to H4 as an internal standard. Error bars denote ±standard deviation.

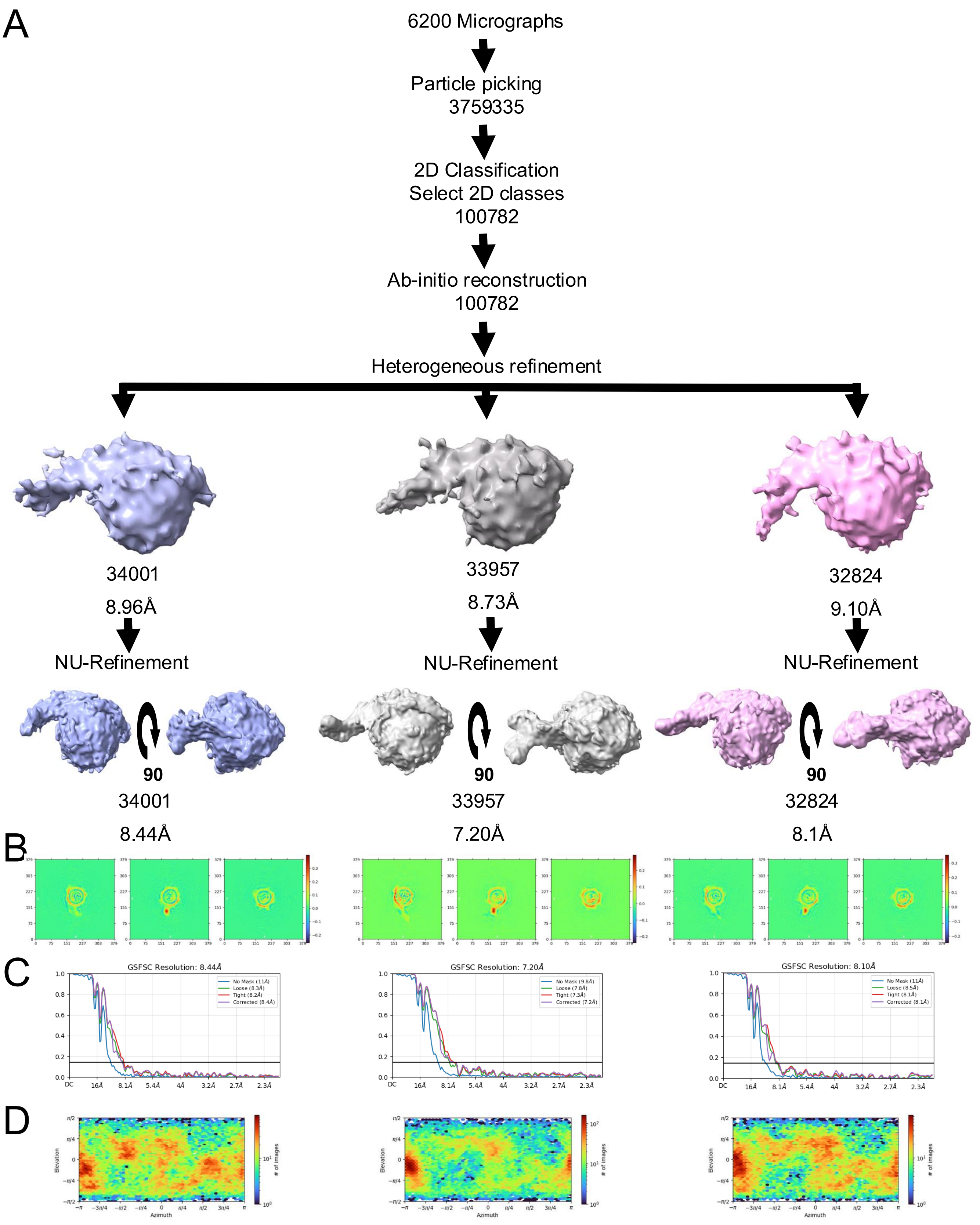


**Figure S6. Workflow for cyro-EM analysis.** *A*, Schematics for cryoEM data analysis using CryoSPARC. The number of the particles and resolution are shown under each group. *B*, Real space slices are shown for the corresponding NU refinement. *C*, GSFSC resolution plots are shown for the corresponding NU refinement. *D*, viewing direction distributions are shown for the corresponding NU refinement.

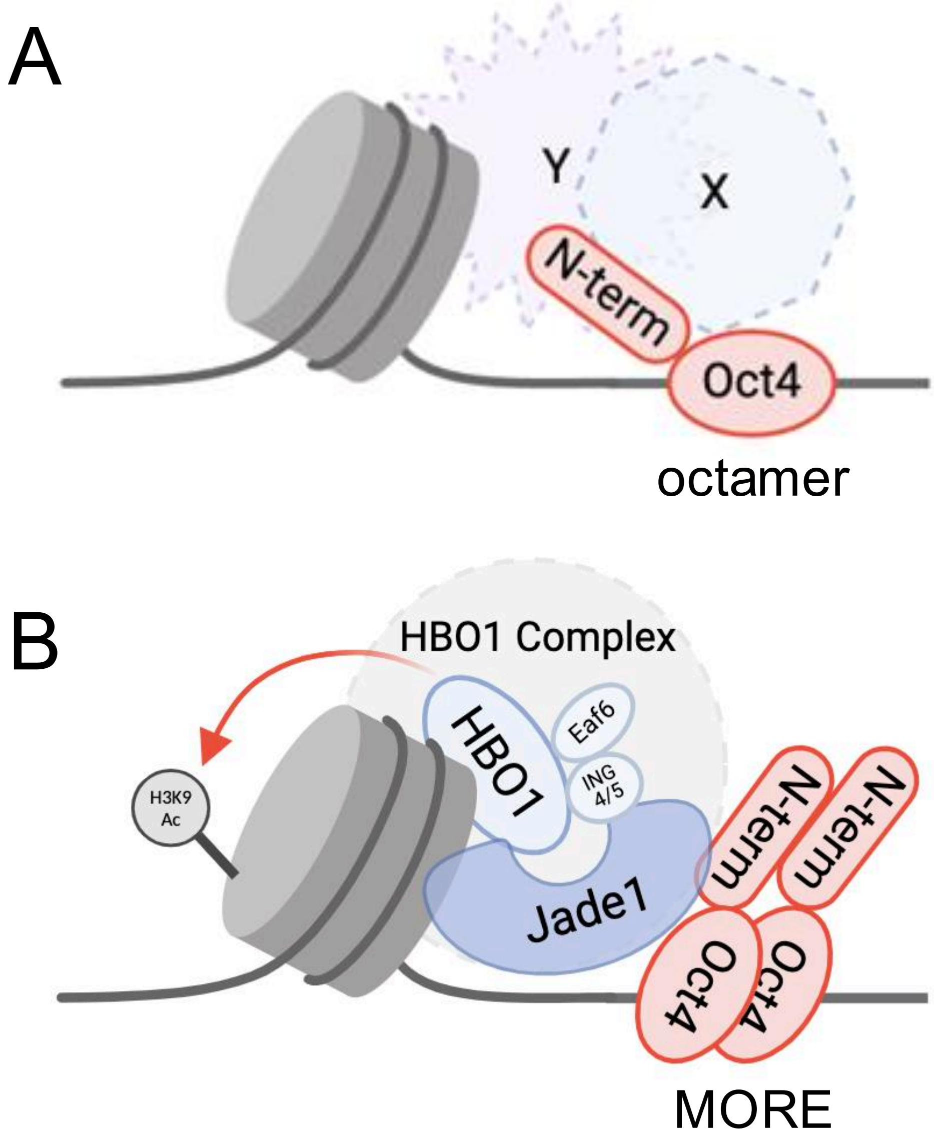


**Figure S7. Model for spatioselective recruitment of Jade1 and HBO1 by multimeric Oct4 at nucleosomes to acetylate H3K9.** *A*, At typical binding sides, Oct4 recruits more generic cofactors (X and Y) to regulate transcription either positively or negatively. *B*, At MORE sites, dimeric Oct4 associates with Jade1 and the HBO1 HAT complex to acetylate H3K9 at local nucleosomes. Other components of the complex such as Eaf6 and ING4 are also shown.
